# SIMON: open-source knowledge discovery platform

**DOI:** 10.1101/2020.08.16.252767

**Authors:** Adriana Tomic, Ivan Tomic, Levi Waldron, Ludwig Geistlinger, Max Kuhn, Rachel L. Spreng, Lindsay C. Dahora, Kelly E. Seaton, Georgia Tomaras, Jennifer Hill, Niharika A. Duggal, Ross D. Pollock, Norman R. Lazarus, Stephen D.R. Harridge, Janet M. Lord, Purvesh Khatri, Andrew J. Pollard, Mark M. Davis

## Abstract

Data analysis and knowledge discovery has become more and more important in biology and medicine with the increasing complexity of the biological datasets, but necessarily sophisticated programming skills and in-depth understanding of algorithms needed pose barriers to most biologists and clinicians to perform such research. We have developed a modular open-source software SIMON to facilitate the application of 180+ state-of-the-art machine learning algorithms to high-dimensional biomedical data. With an easy to use graphical user interface, standardized pipelines, automated approach for machine learning and other statistical analysis methods, SIMON helps to identify optimal algorithms and provides a resource that empowers non-technical and technical researchers to identify crucial patterns in biomedical data.

## Main text

Over the past several years, due to the technological breakthroughs in genome sequencing^1^, high-dimensional flow cytometry^2–4^, mass cytometry^5,6,^ and multi-parameter microscopy^7,8^ the amount and complexity of biological data has become increasingly intractable and it is no longer feasible to extract knowledge without using sophisticated computer algorithms. Therefore, researchers are in need of novel computational approaches that can cope with complexity and heterogeneity of data in an objective and unbiased way. Machine learning (ML), a subset of artificial intelligence, is a computational approach developed to identify patterns from the data in order to make predictions on new data^9^. ML has a profound impact on biological research^10–12^, including genomics^13^, proteomics^14–16^, cell image analysis^17^, drug discovery and development^18^, and cell phenotyping^6,19,20^ which revolutionized our understanding of biological complexity. Recently, using systems-level analysis of genetic, transcriptional, and proteomic signatures to predict patients’ response to vaccines^21,22^, therapies and disease progression^23–27^, ML has become primary computational approach used in the ‘precision medicine’^28^.

The biggest challenge is the proper application of ML methods and the translation of the results into meaningful insights. The analysis of massive datasets and extraction of knowledge using ML requires knowledge of many different computational libraries for data pre-processing and cleaning, data partitioning, model building and tuning, evaluation of the performance for the model and minimizing overfitting^11^. Tools to achieve these tasks have been mainly developed either in R (https://www.r-project.org/)^29,30^ or Python (www.python.org/)^31^, which have today become leading statistical programming languages in data science. Because R and Python are free and open-source, they have been quickly adopted by a large community of programmers who are building new libraries and improving existing ones. As of May 2020, there are 15,658 R packages available in the CRAN package repository (https://cran.r-project.org/). Many of the packages offer different modeling functions and have different syntax for model training, predictions and determination of variable importance. Due to the lack of a unified method for proper application of ML process, even experienced bioinformaticians struggle with these time-consuming ML tasks. To provide a uniform interface and standardize the process of building predictive models, ML libraries were developed, for example mlr3^32^ (https://mlr3.mlr-org.com), the **c**lassification **a**nd **re**gression **t**raining (caret)^30,33^ (https://rdrr.io/cran/caret), scikit-learn^34^ (https://scikit-learn.org), mlPy^35^ (https://mlpy.fbk.eu), SciPy (https://www.scipy.org/) including also ones for deep learning, such as TensorFlow (https://www.tensorflow.org/), PyTorch (https://pytorch.org/) and Keras (https://keras.io/). Since those libraries do not have a graphical user interface, usage requires extensive programming experience and general knowledge of R or Python making it inaccessible for many life science researchers. Therefore, there is an increased effort to harmonize those libraries and develop a software that will facilitate application of ML in life sciences.

The software should provide a standardized ML method for data pre-processing, data partitioning, building predictive models, evaluation of model performance and selection of features. Moreover, such software should be adapted for biological datasets that have high percentage of missing values^36^, have imbalanced participant distributions (i.e. having a high number of infected subjects, but only a relatively small number of healthy controls)^37^ or suffer from a *“curse of dimensionality”*, i.e. poor predictive power, as can be observed when the number of features is much greater than the number of samples^38^. Additionally, beyond ML process, the software should support exploratory analysis and visualization of the results using user-friendly graphical interface. The fast-paced technological development dramatically increased size of biological datasets and computational power needed for analysis. Therefore, open-source web-based software supporting cloud processing architecture is essential. Additionally, software should support an automated ML^39^ (autoML) process that rapidly builds high-performance predictive models by identifying optimal ML method, including selection of an appropriate algorithm, optimization of model hyperparameters and evaluation of the best-performing models. AutoML improves the efficiency of ML process and resulting models often outperform hand-designed models^39,40^.

To address these challenges, we developed SIMON (Sequential Iterative Modeling “Over Night”), a free and open-source software for application of ML in life sciences that facilitates production of high-performing ML models and allows researchers to focus on knowledge discovery process. SIMON provides a user-friendly, uniform interface for building and evaluating predictive models using a variety of ML algorithms. Currently, there are 182 different ML algorithms available (**Supplementary table 1**). The entire ML process which is based on the caret^33^ library, from model building and evaluation to feature selection in SIMON is fully automated. This allows advanced ML users to focus on other important aspects necessary to build highly accurate models, such as data preprocessing, feature engineering and model deployment. It also makes the entire ML process more accessible to domain-knowledge experts that formulated the research hypothesis and collected the data, but lack programming ML skills Additionally, to prevent optimistic accuracy estimates and to optimize the model for generalization to unseen data, SIMON introduces unified process for model training, hyperparameter tuning and model evaluation by generation of training, validation and test sets. Training set is used for building models, validation set is used for hyperparameter tuning and finally, models are evaluated in an unbiased way using the test, also known as holdout set that has never been used in training. Beside the standardized ML process, the initial install version offers a set of core components specifically suited for analysis of biomedical data, such as multi-set intersection function for integration of data with many missing values^41^ (https://cran.r-project.org/web/packages/mulset/index.html), method for identifying differentially expressed genes using significance analysis in microarrays (SAM)^42^, graphical representation of the clustering analysis important for detection of batch effects, graphical display of the correlation analysis and graphical visualizations of the ML results that can be downloaded as publication-ready figures in scalable vector graphics (SVG) format. Finally, SIMON is available in two versions as a single mode and a server version. The single mode is developed as a SIMON Docker container (https://www.docker.com/), ensuring code reproducibility and solving installation compatibility issues across major operating systems (Windows, MacOS and Linux). In both versions parallel computing is supported which is essential for more efficient ML analysis by distributing the workload across several processors. To promote collaboration, data sharing and support distributed cloud processing, SIMON is also available as a server version. The server version can be installed on a private or a public Linux cloud service. Distributed cloud processing (multiNode) is implemented utilizing OpenStack, a free and open source cloud computing platform (https://www.openstack.org/). The advantage of the server versions is that it has multiNode capability which allows users to distribute workload on multiple computers simultaneously to optimize SIMON performance. The multiNode process can be used to horizontally scale analysis to large infrastructure, such as high performance computing clusters to meet the computational needs and accommodate parallel processing of large amounts of data. Additionally, in the server version, users can configure data storage either on a local server or in a cloud using service which is interoperable with Amazon Web Services S3 application programming interface (AWS S3 API)^43^. SIMON is also translated into multiple languages by collaborative open-source effort. SIMON source code is regularly updated, and both source code and compiled software are available from the project’s website at http://www.genular.org/. Overall, SIMON is designed to provide a uniform knowledge discovery interface adaptable to the increasing size of biomedical datasets allowing data scientists, bioinformaticians and domain-knowledge experts to solve biological research questions.

We demonstrate the accuracy, ease of use and power of SIMON on five different biomedical datasets and build predictive models for arboviral infection severity (SISA)^44^, the identification of the cellular immune signature associated with a high-level of physical activity (Cyclists)^45^, the determination of the humoral responses that mediate protection against *Salmonella* Typhi infection (VAST)^46^, early-stage detection of colorectal cancer from microbiome data (Zeller)^47,48,^ and for the detection of liver hepatocellular carcinoma cells (LIHC)^49^ (**Fig. 1 b, c, d, e, Supplementary protocol**). To build models using the SISA dataset (described in the **Supplementary methods** and available as **Supplementary table 2**), 11 ML algorithms were used, five from the original publication^44^ (treebag, k nearest neighbors, random forest, stochastic generalized boosting model and neural network) and additionally, ‘*sda’,* shrinkage discriminant analysis; *‘hdda’,* high dimensional discriminant analysis; *‘svmLinear2’*, support vector machine with linear kernel; *‘pcaNNet’*, neural networks with feature extraction; *‘LogitBoost’,* boosted logistic regression and naïve Bayes. Due to the unified ML process for training, tuning and evaluating predictive models, users can test a variety of ML algorithms in SIMON. Since the same training and test sets are used by different algorithms, resulting models can be compared and the best performing models can be selected. After manually setting initial parameters for data partitioning, predictor and outcome variables, exploratory classes, pre-processing and selecting ML algorithms (**Fig. 1a**), SIMON automatically performs all necessary ML analysis steps to build, tune and evaluate predictive models. The process of building all 11 models on the SISA dataset in SIMON finished in 59 sec on a standard laptop (Intel® Core™ i7 Processor 7700HQ and 16 GB of RAM). In SIMON, users can evaluate model performance using standard performance measurements such as accuracy, sensitivity, specificity, precision, recall, area under the receiver operating characteristic curve (AUROC), precision-recall area under curve (prAUC), and logarithmic loss (LogLoss) on training and holdout, test sets (**Fig. 1b**). The shrinkage discriminant analysis model (‘*sda’*) had the highest AUROC of 0.97 on the training set and also performed well on the holdout, test set (test AUROC 0.96) (**Fig. 1c, Supplementary table 3,** the model is available as the **Supplementary data 1**).

**Figure 1.**
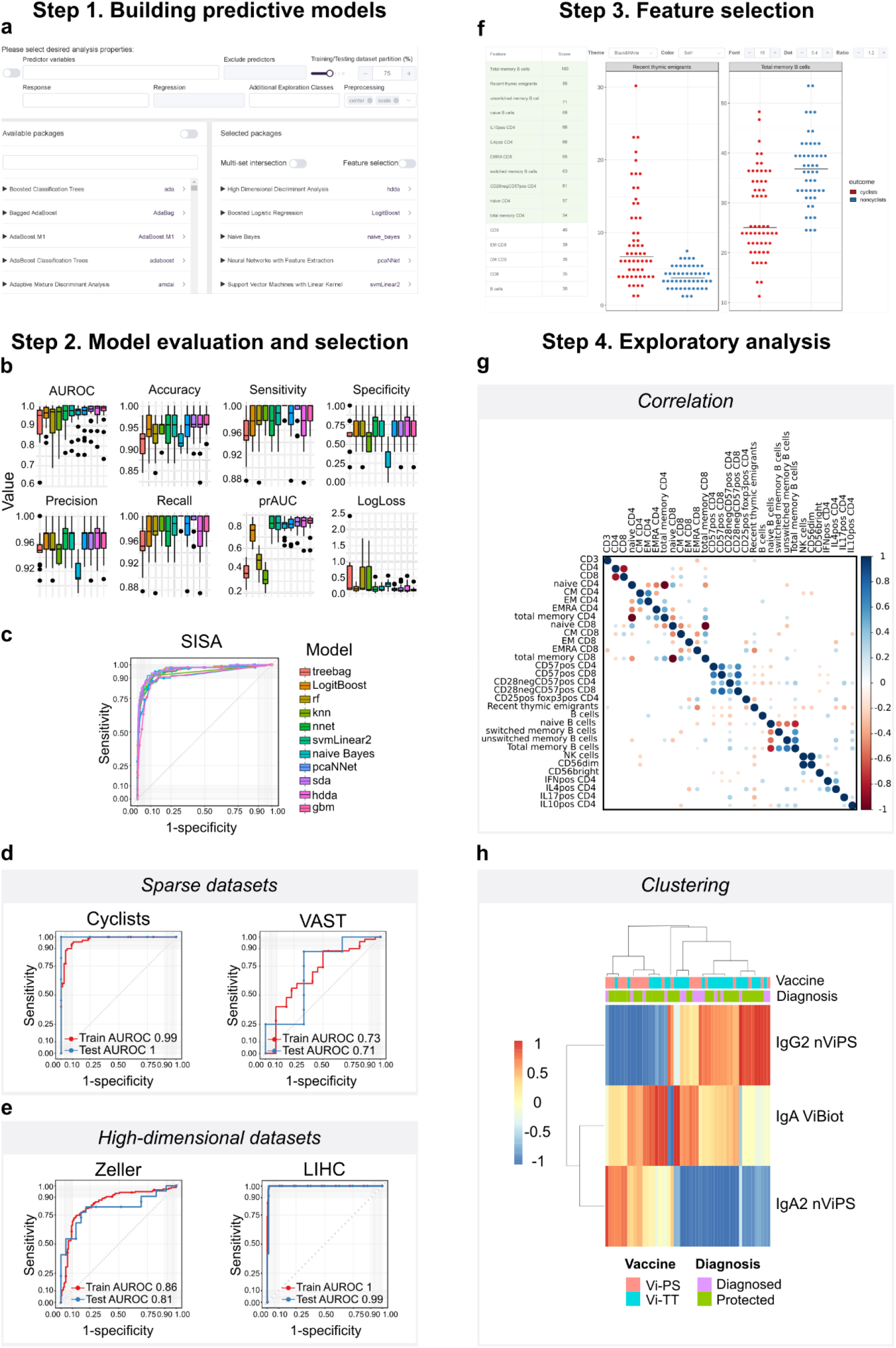
SIMON machine learning workflow. Step 1. Building predictive models. **(a)** Screenshot of the SIMON graphical user interface demonstrating input selection for machine learning analysis, such as predictors and response (outcome) variables, additional exploration classes, training/test split, preprocessing functions and desired machine learning algorithms. **Step 2. Model evaluation and selection.** Comparison of **(b)** box plots of performance measurements calculated for 11 predictive models and **(c)** ROC curves built on the SISA dataset. Comparison of ROC curves calculated from the training and test sets on **(d)** datasets with missing values (Cyclists and VAST) and **(e)** high-dimensional datasets (Zeller and LIHC). **Step 3. Feature selection. (f)** Screenshot of the SIMON interface showing variable importance scores calculated for each feature and graphical visualization of the selected features from the Cyclists dataset. **Step 4. Exploratory analysis. (g)** Correlation analysis on the Cyclists dataset. **(h)** Clustering analysis on the VAST dataset.

To demonstrate SIMON’s capabilities for analyzing biomedical datasets with missing data, we applied SIMON to (i) the Cyclists dataset studying the impact of physical activity on the immune system in adulthood^45^ (**Supplementary table 4**) and (ii) the VAST dataset collected from a clinical trial which was undertaken to evaluate typhoid vaccine efficacy^50^ (**Supplementary table 5**). Description of both datasets is available in **Supplementary methods.** The percentage of missing values was 8% in the Cyclists dataset and 21% in the VAST dataset either due to the exclusion of samples not passing quality control criteria or the lack of sample volume to repeat experiments and obtain reportable data. To build models using the datasets with missing values, we used the multi-set intersection (*mulset*) function^41^ to identify shared features between donors and generate resamples (**Supplementary protocol**). Because *mulset* function generates multiple resamples from the initial dataset based on shared features, it is useful for removal of missing values and can be used for integration of data collected from different assays and across clinical studies^41^. For the Cyclists dataset, the *mulset* function generated 146 resamples. The models were built for each of the 146 resamples using five ML algorithms (naïve Bayes, svmLinear2, pcaNNet, logistic regression and hdda) to identify immune cell subsets enriched in the cohort of master cyclists. The analysis finished in 41 min and 24 sec. The model with the highest performance measures was built with naïve Bayes on the resample with 96 donors that shared 31 features (train AUROC 0.99 and test AUROC 1) (**Fig. 1d, Supplementary table 6 and Supplementary data 2**). The *mulset* function generated 206 resamples from the initial VAST dataset with varying number of donors and features. Resamples with less than 10 donors in the test set were removed prior ML process to prevent too optimistic predictive estimates using the holdout set. Therefore, the ML analysis was performed on 58 resamples using same five ML algorithms as for the Cyclists dataset. The entire analysis finished in 31 min and 1 sec. The top performing model was built on the resample with 47 donors that shared 13 features with the naïve Bayes algorithm (train AUROC 0.73 and test AUROC 0.71) (**Fig. 1d**, **Supplementary table 7 and Supplementary data 3**).

We also applied SIMON to (i) a dataset with a large number of features measured using whole-metagenome shotgun sequencing of fecal samples (Zeller dataset, **Supplementary table 8**), and (ii) the liver hepatocellular carcinoma dataset from TCGA with an imbalanced sample distribution of tumor and adjacent normal tissue samples (LIHC dataset, **Supplementary table 9**). Both datasets are described in **Supplementary methods**. For the Zeller dataset, models were built using ML algorithms known to perform well in the situations where more features were measured than individuals, such as shrinkage discriminant analysis^51^, high dimensional discriminant analysis^52^ and neural network with feature extraction^53^. Two additional algorithms were included, svmLinear2 and LogitBoost. The complete analysis was performed in less than 1 min (0:38 min). The sda algorithm built the model with the highest performance (train AUROC 0.86 and test AUROC 0.81) having a higher performance measure than the published LASSO linear regression model^47^ (train AUROC 0.84 and test AUROC 0.85) (**Fig. 1e, Supplementary table 10 and Supplementary data 4**). For the LIHC dataset we used same five ML algorithms as for Zeller dataset and analysis finished in 11 min and 30 sec. For such highly imbalanced dataset the precision-recall AUC (prAUC)^54^ is a much better performance measurement than AUROC that reported near-perfect performance (**Fig. 1e**). The prAUC provides information how well the model correctly detects cancer cells, while it is less stringent on the evaluation of healthy cells. To avoid obtaining overly optimistic prediction results (often observed on imbalanced datasets), we ranked models based on the prAUC of the training set (**Supplementary table 11)**. The model that had the best performance was built using the svmLinear2 algorithm (train prAUC 0.83) and it also performed well on the holdout, test set (prAUC 0.73) (**Supplementary data 5**).

The *drowsiness* contributed the most to the top-performing SISA model, confirming the findings from the original study^44^ (**Supplementary table 12**). To standardize the process for evaluation of the features and their contribution to the models, we implemented the variable importance score evaluation functions from the caret library^33^. This allows users to compare features selected across models. In the case of SISA dataset, *drowsiness* contributed the most in all of the models built (**Supplementary table 13**), indicating the importance of this symptom and its correlation with hospitalization. The features that contributed the most to the Cyclists model were total memory, unswitched memory and naïve B cells, recent thymic emigrants, CD8+ T cells with TEMRA phenotype, and regulatory T cells (CD25+ Foxp3+ CD4+ T cells) (**Supplementary table 14**). In comparison to age-matched physically inactive individuals (non-cyclists), the master cyclists had increased frequencies of recent thymic emigrants, naïve B cells and CD3 cells, and decreased frequencies of memory B cells and CD8 T cells with TEMRA phenotype, confirming that ageing of immune systems i.e. immunosenescence can be reduced by high levels of physical activity^45^ (**Fig. 1f, Supplementary figure 1**). To further explore the relationship between selected features, users can perform correlation analysis to reveal highly correlated features (**Fig. 1g, Supplementary protocol**). Naïve and memory B cells were identified as being highly correlated (**Fig. 1g**), as expected since these subsets were determined from the same flow cytometry plots and their relationship is inversely correlated. Removal of those highly correlated features can help to build more accurate models. Removal of naïve B cells resulted in building predictive model with the same performance measurements as the model built on the entire dataset (train AUROC 0.99 and test AUROC 1) (**Supplementary table 15**), while removal of total memory B cells lowered the accuracy estimates (train AUROC 0.98 and test AUROC 1) (**Supplementary table 16**), indicating the importance of memory B cells to discriminate between master cyclists and non-cyclists. In the VAST dataset, individuals with higher IgA, IgA1, IgA2 and IgG2 titers against native Vi polysaccharide (nViPS) antigen and higher IgA and IgG3 titers against biotinylated Vi polysaccharide (ViBiot) on the day of the challenge were protected against the typhoid challenge supporting the data from univariate analysis^46^ (**Supplementary table 17 and Supplementary figure 2**). Moreover, using the clustering function of SIMON’s exploratory analysis module, we can quickly identify that the IgA2 signature dominates the responses after vaccination with a purified Vi polysaccharide (Vi-PS), while the IgG2 signature was dominant for the Vi tetanus toxoid conjugate (Vi-TT) vaccine^46^ (**Fig. 1h, Supplementary protocol**). For the Zeller dataset, the same features as originally reported^40^ contributed the most to the model, including *Fusobacterium nucleatum* and *Peptostreptococcus stomatis* (**Supplementary table 18**). The features that contributed the most to the LIHC model were well-known genes identified to be upregulated in LIHC such as *GABRD* and *PLVA*P^*55*^ and genes enriched in adjacent normal tissue samples *ANGPTL6*^*56*^, *VIPR1^57^* and *OIT3*^*58*^ as a typical signature for healthy liver tissue (**Supplementary table 19**, **Supplementary figure 3**).

Overall, SIMON is a powerful software platform for data mining that facilitates pattern recognition and knowledge extraction from high-quality, heterogenous biological and clinical data, especially where there is missing data, an imbalanced distribution and/or high dimensionality. It can be used for identification of genetic, microbial and immunological correlates of protection and help guiding further analysis of the biomedical data.

