## Supplementary figures and images for "SIMON: open-source knowledge discovery platform"

### Supplementary Figure 1

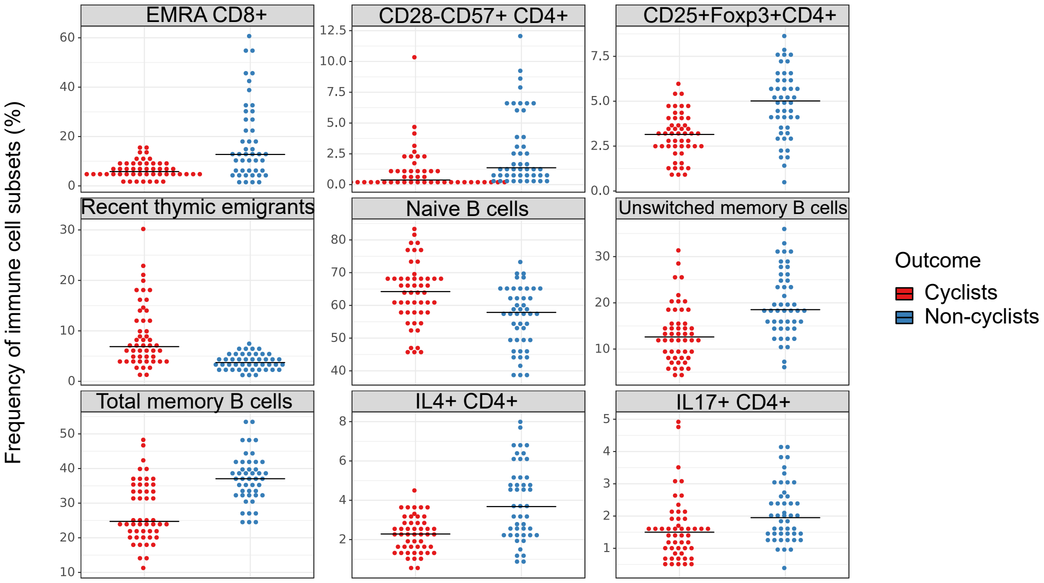


Tomic et al. Supplementary Figure 1

### Supplementary Figure 2

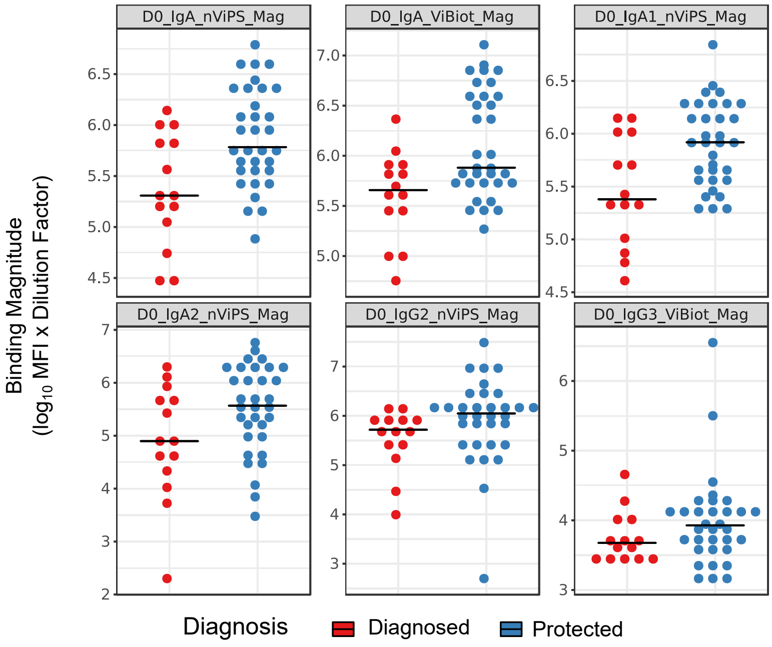


Tomic et al. Supplementary Figure 2

### Supplementary Figure 3

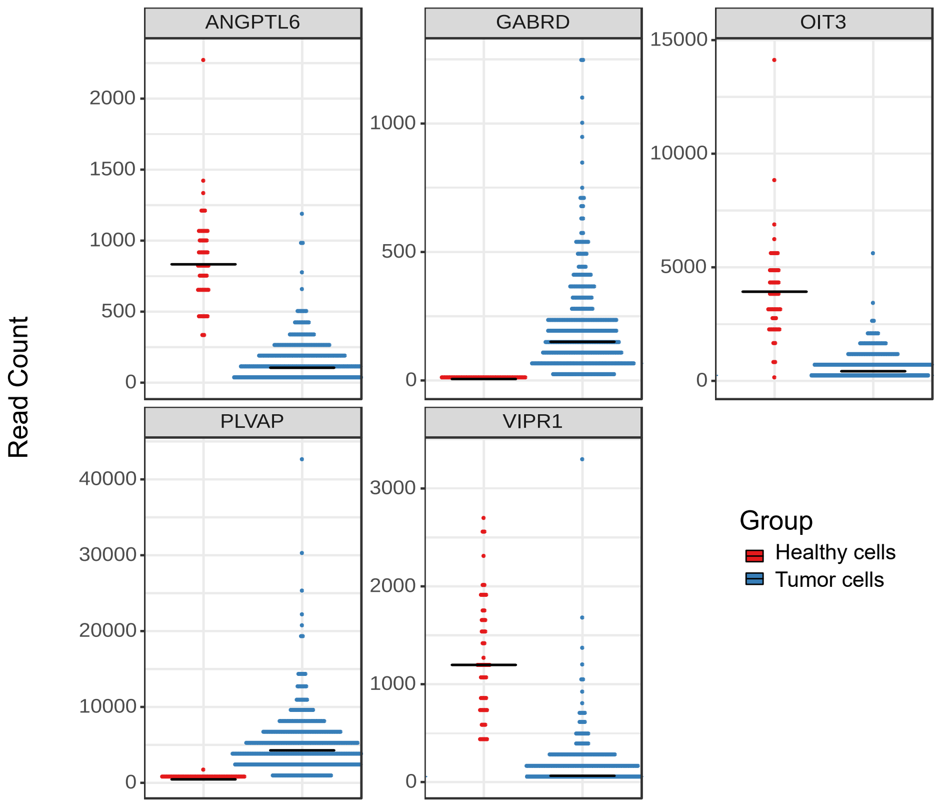


Tomic et al. Supplementary Figure 3
