## Supplementary Materials for "SIMON: open-source knowledge discovery platform"

### SIMON: open-source machine learning knowledge discovery platform

[1]Oxford Vaccine Group, Department of Pediatrics, University of Oxford, Oxford, UK. [2]Institute of Immunity, Transplantation and Infection, Stanford University School of Medicine, Stanford, CA, USA. [3]Deep Medicine, Nuffield Department of Women's and Reproductive Health, University of Oxford, Oxford, UK. [4]Graduate School of Public Health and Health Policy and [5]Institute for Implementation Science and Population Health, City University of New York, NY, USA. [6]RStudio, PBC, Boston, MA, USA. [7]Duke Human Vaccine Institute, Duke University, Durham, NC, USA. [8]MRC-Versus Arthritis Centre for Musculoskeletal Ageing Research, Institute of Inflammation and Ageing, University of Birmingham Research Labs, Birmingham, UK. [9]Centre for Human and Applied Physiological Sciences, King's College London, UK. [10]NIHR Birmingham Biomedical Research Centre, University Hospital Birmingham NHS Foundation Trust and University of Birmingham, UK. [11]Center for Biomedical Informatics Research, Department of Medicine, Stanford University, Stanford, CA, USA. [12]Department of Microbiology and Immunology, Stanford University School of Medicine, Stanford, CA, USA. [13]Howard Hughes Medical Institute, Stanford University, Stanford, CA, USA.

#### Supplementary methods

**Datasets.** Datasets used in the **Figure 1** were obtained either directly from authors (VAST[51] and Cyclists[50] datasets) or downloaded from publication[49] (SISA dataset) and R packages (Zeller dataset from the *MetagenomicData*[53] and LIHC from the *GSEABenchmarkeR*[54]) with the help from the authors. The SISA dataset contains data from 543 individuals hospitalized due to the arboviral infection with Dengue, chikungunya or Zika viruses from a surveillance study in Ecuador collected from 2013 to 2017. In the SISA dataset we have excluded columns with high level of missing values (pregnancy, *WomPreg* and complete blood count test which was not performed for all donors and includes columns: *PLT_count*, *Lymphocytes*, *CBC_N%*, *WBC_calc* and *CBC_HCT*). Additionally, 9 donors with missing values were removed. The final SISA dataset after removal of columns and row with missing values is available as **Supplementary table 2**. The Cyclists dataset contains data from immune responses of 120 elderly individuals with the high-level of physical activity i.e. master cyclists and 75 age-matched controls with low level of physical activity (non-cyclists) analyzed using flow cytometry (**Supplementary table 4**). The VAST dataset contains data from 72 individuals enrolled in the clinical study to evaluate humoral responses in a typhoid vaccine efficacy trial in a controlled human infection model. Only day 0 (day of the challenge) log-transformed data were used and are available for download as **Supplementary table 5**. Individuals were vaccinated with either a purified Vi polysaccharide (Vi-PS) vaccine (35 individuals) or the Vi tetanus toxoid conjugate (Vi-TT) vaccine (37 individuals) one month prior to oral challenge with live *Salmonella* Typhi. Out of 72 individuals, 26 developed an acute typhoid infection following challenge. The Zeller dataset contains information on the microbiome species abundance in healthy individuals and colorectal cancer patients (**Supplementary table 8**). The data was access through *MetagenomicData* package. In total 184 individuals were included of which 93 were healthy controls and 91 colorectal cancer patients. The LIHC dataset obtained from the *GSEABenchmarkeR* package contains RNA expression data from 374 liver hepatocellular carcinoma (LIHC) cells and 50 adjacent normal cells (**Supplementary table 9**).

**Installing SIMON.** SIMON can be installed directly from the GitHub (https://github.com/genular/simon-frontend) or a pre-built version can be installed from DockerHub (https://www.docker.com/). Users need to install Docker (version 17.05 or later required) following instructions available on the Docker website (https://docs.docker.com/). Installation instructions for Windows (https://docs.docker.com/docker-for-windows/install/), MacOS (https://docs.docker.com/docker-for-mac/install/) and Linux (https://docs.docker.com/install/linux/docker-ce/ubuntu/) are provided. After Docker installation, users must download and run a SIMON image from DockerHub. To do that users must run Terminal on Linux and MacOS or Windows Power Shell if using Windows OS and type following command:

```
docker run --rm --detach --name genular --tty --interactive --env
IS_DOCKER='true' --env TZ=Europe/London --volume
genular_data:/mnt/usrdata --publish 3010:3010 --publish 3011:3011 --
publish 3012:3012 --publish 3013:3013 genular/simon:latest
```

Variable 'TZ=' stands for time zone and can be replaced with appropriate time zone. Once command is executed, SIMON will be downloaded and started. To access SIMON open web browser (Firefox recommended, available at https://www.mozilla.org/firefox/) and type: `http://localhost:3010`. Create administrator account. SIMON will run until you shutdown/restart your computer or stop it manually using following command: `docker stop genular`. Advance instructions for installing a server version are provided on our GitHub page.

**Code and data availability.** The source code of the SIMON is available at https://github.com/genular/simon-frontend. All data used in SIMON analysis are available as Supplementary tables, while models built are available as Supplementary data in the RData format.

#### Supplementary Figures, Tables and Data

Fig. S1. Frequency of immune cell subsets associated with high-level of physical activity.

Fig. S2. Antibody-mediated signature associated with the effective vaccine against *Salmonella* Typhi infection.

Fig. S3. Gene expression signature specific for tumor cells.

Table S1. List of machine learning algorithms implemented in SIMON.

Table S2. SISA dataset.

Table S3. List of models built for the SISA dataset.

Table S4. Cyclists dataset.

Table S5. VAST dataset.

Table S6. List of models built for the Cyclists dataset.

Table S7. List of models built for the VAST dataset.

Table S8. Zeller dataset.

Table S9. LIHC dataset.

Table S10. List of models built for the Zeller dataset.

Table S11. List of models built for the LIHC dataset.

Table S12. List of features and their variable importance score in the SISA dataset selected in the top performing model.

Table S13. List of features and their variable importance score in the SISA dataset selected in all models.

Table S14. List of features and their variable importance score in the Cyclists dataset.

Table S15. List of models built after removal of naive B cells.

Table S16. List of models built after removal of memory B cells.

Table S17. List of features and their variable importance score in the VAST dataset.

Table S18. List of features and their variable importance score in the Zeller dataset.

Table S19. List of features and their variable importance score in the LIHC dataset.

Data file S1. RData sda model for the SISA dataset.

Data file S2. RData naïve Bayes model for the Cyclists dataset.

Data file S3. RData naïve Bayes model for the VAST dataset.

Data file S4. RData sda model for the Zeller dataset.

Data file S5. RData svmLinear2 model for the LIHC dataset.

#### Supplementary figures

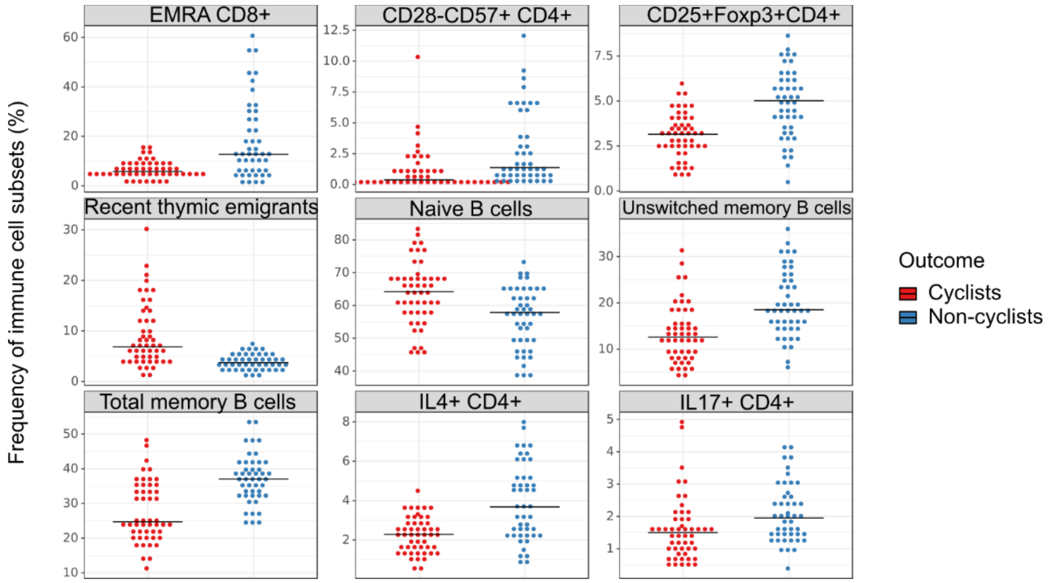


**Supplementary figure 1. Frequency of immune cell subsets associated with high-level of physical activity.** Dot plots represent distribution of immune cell subsets between cyclists (red dots) and non-cyclists (blue dots) as frequency (percentage of parent immune cell population) for the top nine selected features that contribute the most to the Cyclists model to discriminate between cyclists and non-cyclists. Each dot is one individual, lines indicate median values.

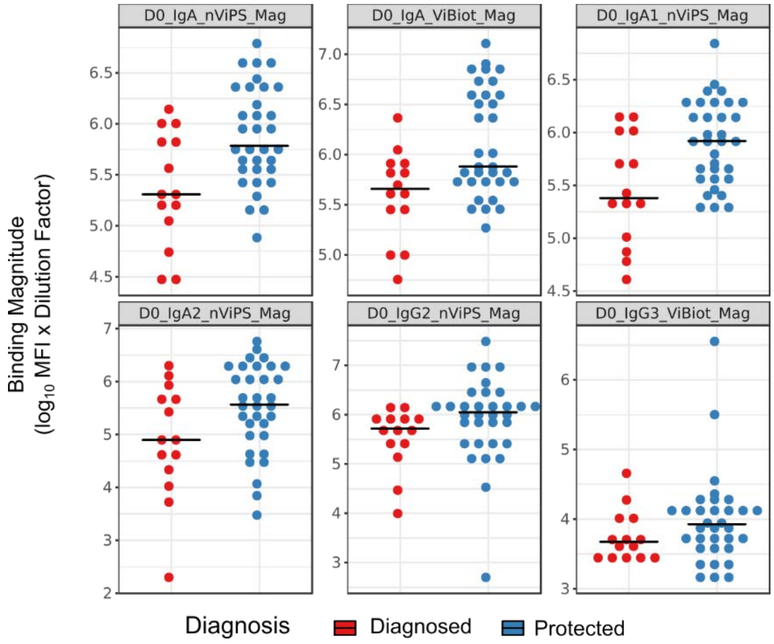


**Supplementary figure 2. Antibody-mediated signature associated with the effective vaccine against *Salmonella* Typhi infection.** Dot plots represent binding magnitude of indicated antibodies between diagnosed (red dots) and protected (blue dots) individuals. Each dot represents one individual, while lines indicate median values. The binding magnitude is log-transformed and given as Mean Fluorescence Intensity (MFI) multiplied by dilution factor. *D0*, day 0 (day of the challenge); *nViPS*, native Vi polysaccharide antigen; *ViBiot*, biotinylated Vi polysaccharide antigen; *Mag*, magnitude.

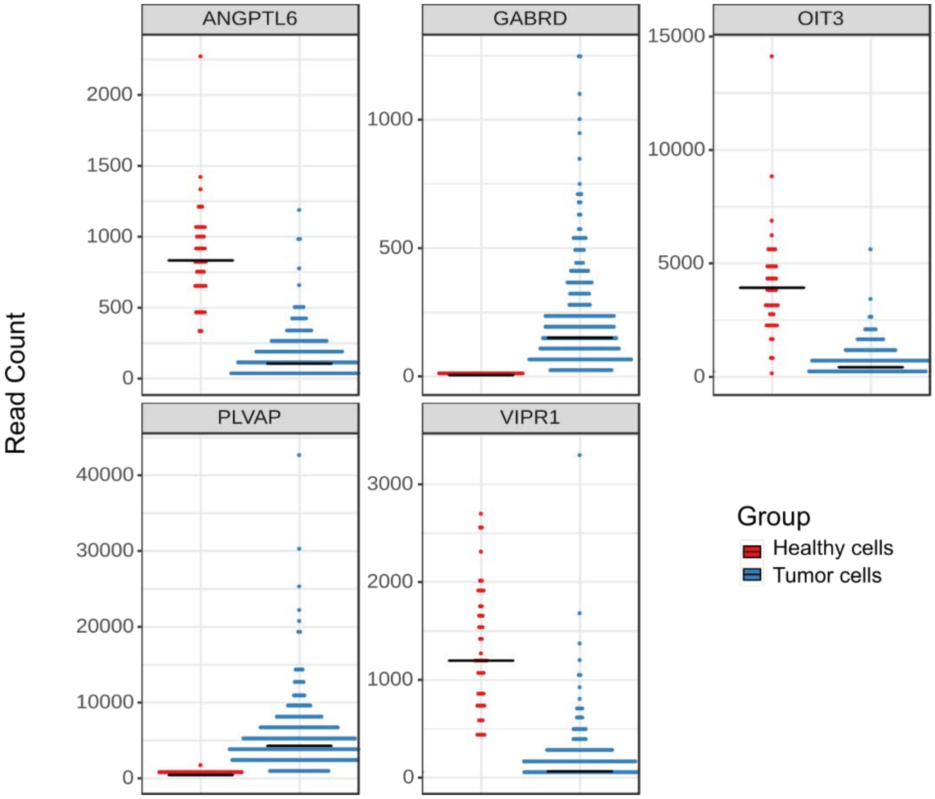


**Supplementary figure 3. Gene expression signature specific for tumor cells.** Dot plots represent read counts for top five genes that discriminate between healthy (red dots) and tumor (blue dots) cells selected in the top performing model. Each dot represents one sample, while lines indicate median values.

#### Supplementary protocol

Step-by-step instructions how to run SIMON analysis for the following use cases: (1) Identifying clinical biomarkers that can predict the severity of the arboviral infection severity; (2) Predicting antibody signature to mediate protection against *Salmonella* Typhi challenge infection; (3) Identifying cellular immune signature associated with high-level of physical activity; (4) Building predictive model for the early-stage detection of colorectal cancer using microbiome; and (5) Building predictive model for detection of liver hepatocellular carcinoma cells using transcriptome data.

**Use case 1. Identifying clinical biomarkers that can predict the severity of the arboviral infection severity.**

**Step 1. Uploading data.** SISA dataset (available as **Supplementary table 2**) needs to be uploaded as CSV file in the following format: donors/samples in rows and features that were measured (i.e. clinical measurements) in columns (*'Click here'* red arrow on image below). One of the columns contains information about the outcome, in this case this is the column named *'Hospitalized'* and the outcome is labelled with zero if *'non-hospitalized'* and with one is *'hospitalized'*. Note that SIMON can analyze data using either text or numeric values for the outcome variable.

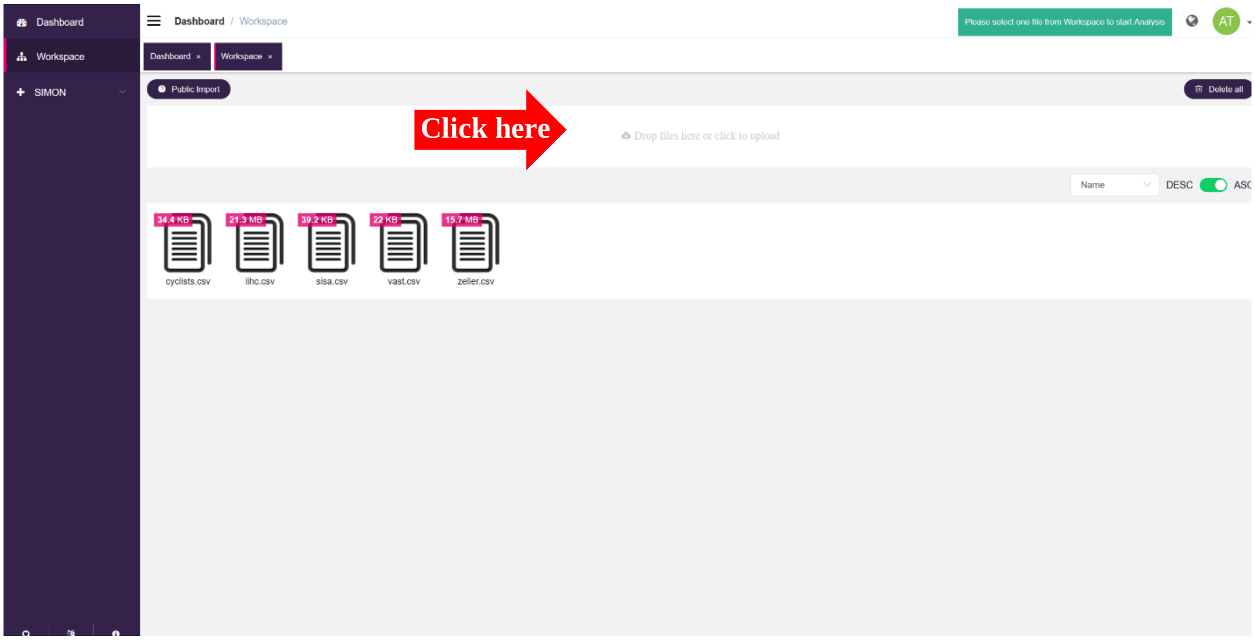


**Step 2. Selecting dataset.** To start the analysis, user must select the SISA dataset by clicking the document icon (A). The selected SISA dataset will be highlighted in grey and in the upper right-hand side the green tab will show the name of the SISA dataset (B). Now in the left-hand side menu (in purple) the *'Analysis'* tab becomes available (C).

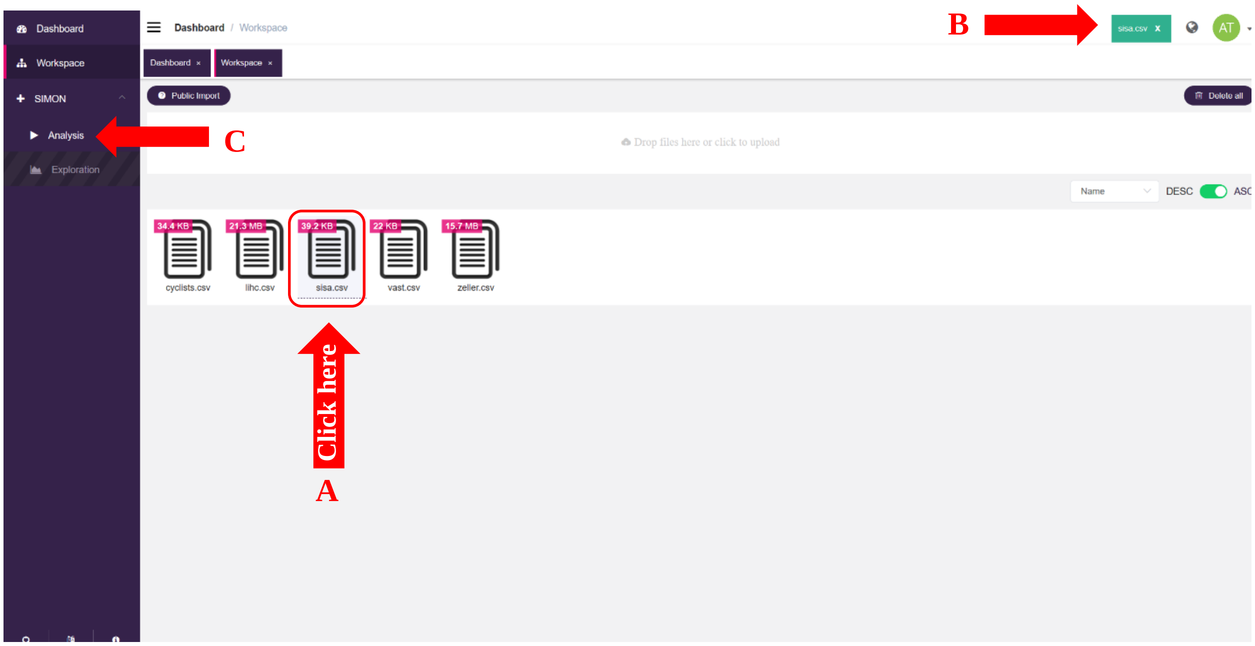


**Step 3. Setting-up analysis.** Once SISA dataset is selected, users can start with the analysis by clicking the '*Analysis*' on the left-hand side menu (in purple). This opens new window '*Analysis*' where users can select analysis parameters and ML algorithms. For the SISA dataset, in the Box 1, users must select all predictor variables by clicking the button next to the input form (A) since we want to use all clinical measurements for the prediction model. Alternatively, if users want to select only some features, by clicking in the input field '*Predictor variables*' first 50 available columns are shown in the drop-down menu and users can choose which columns they want to use for analysis. If there are more than 50 columns available, users can type which columns they want to use. Next, we select the outcome we want to predict in the '*Response*' input field, in the SISA dataset that is the '*Hospitalized*' column (B). We then select which columns to exclude (C). In the SISA dataset we have excluded column without any information for the predictive model (donor identification numbers in the *IDindex* column). The initial SISA dataset is split into training (85% of the data) and test sets (15% of the data) (D). Finally, for the pre-processing step, data was centered (mean subtracted from values) and scaled (values divided by standard deviation) (E).

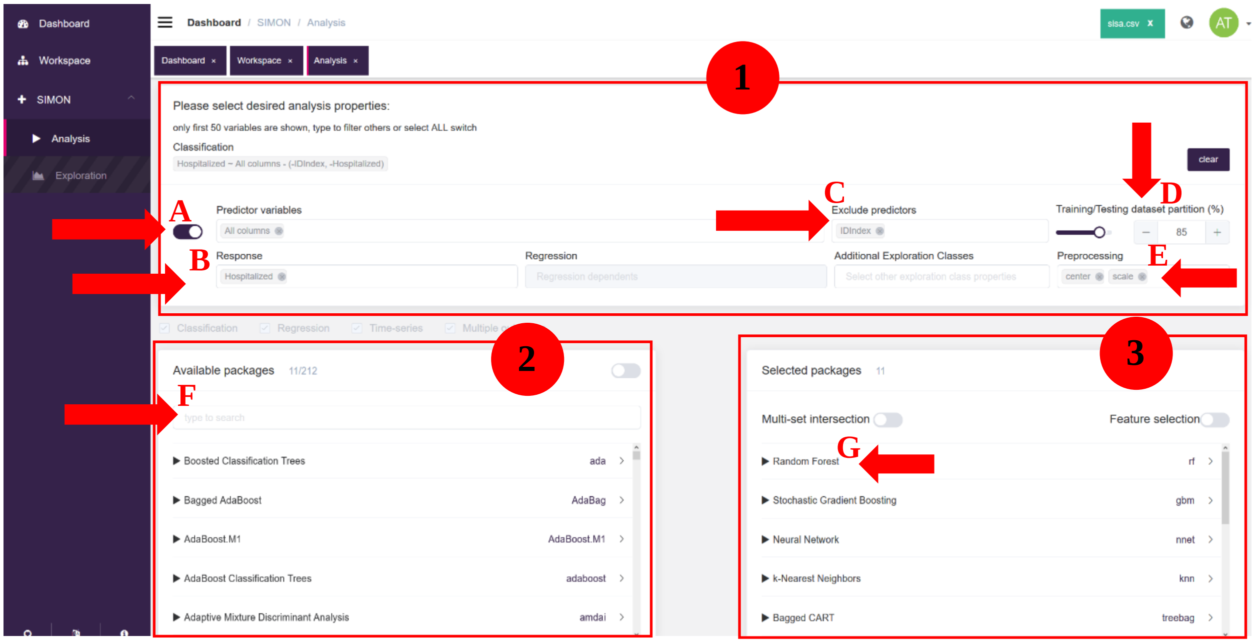


Users can choose which pre-processing functions they want to apply. Available pre-processing functions are Box-Cox transformation (BoxCox), power transformation (expoTrans), Yeo-Johnson transformation (YeoJohnson), subtract mean from values (center), divide by standard deviation (scale), normalize values (range), K-nearest neighbors Imputation (knnimpute), Imputation using Bagging of regression tress (bagImpute), median impute (medianImpute), prinicipal component analysis (pca), data projected onto a unit circle (spatialSign), correlation filtering (corr), remove zero-variance (zv), remove near zero-variance (nzv) and exclude predictors that have only one unique value (conditional).

In the Box 2, users select which ML algorithms to use, while 5 of the default ML algorithms are already selected in the Box 3. For the analysis of the SISA dataset, we will, in addition to five already selected, select additional six ML algorithms: shrinkage discriminant analysis, treebag, k nearest neighbors, random forest, stochastic generalized boosting model and neural network. Name of the ML algorithm is typed in the input field (F). The full names of the packages for the selected algorithms are: *'Shrinkage discriminant analysis'*, Shrinkage discriminant analysis (sda); *'Treebag'*, Bagged CART; *'k nearest neighbors'*, k-Nearest Neighbors (knn); *'Random forest'*, Random forest (rf); *'Stochastic generalized boosting model'*, Stochastic gradient boosting (gbm) and *'Neural network'*, Neural network (nnet). Once the name of the algorithm is typed and user clicks on the desired package, that algorithm is automatically added to the list of selected algorithms in the Box 3. Note, that sometimes different R packages are available for same algorithm, as it is the case for Random forest algorithm. SIMON allows users to inspect selected algorithms by clicking on their names (G). Users then obtain additional information about the algorithm (H) and they can click to obtain the reference to the original publication (I) to be sure that they select appropriate algorithms.

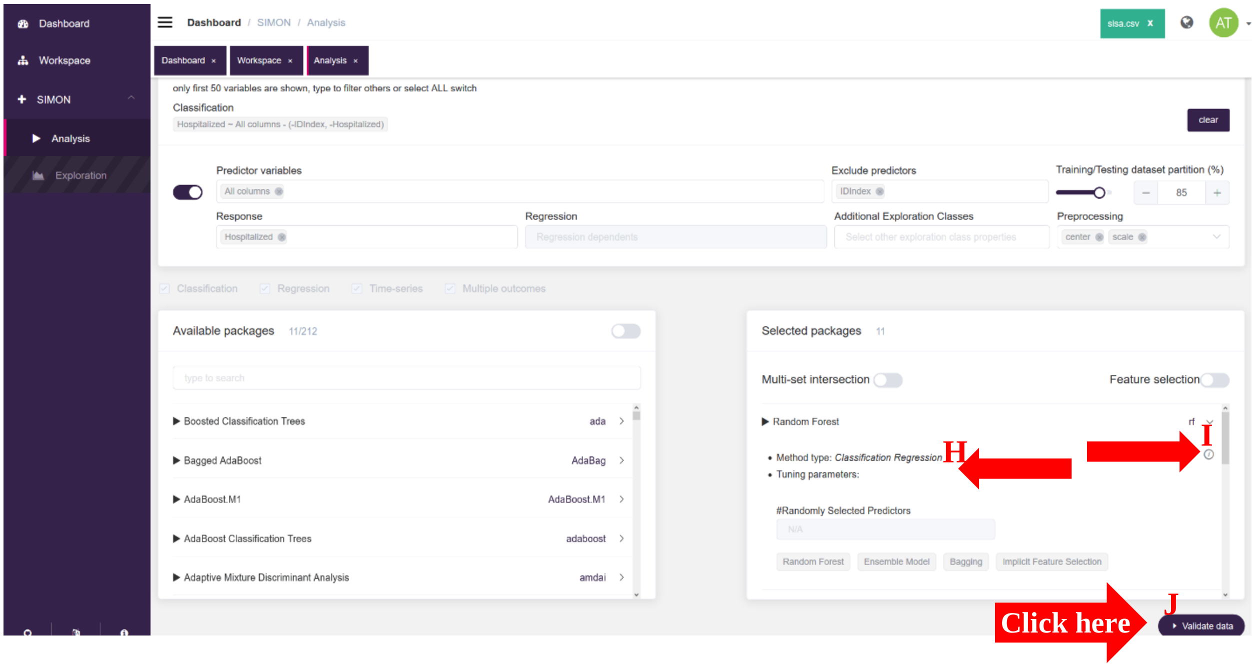


Finally, analysis is initiated by clicking the '*Validate data*' button (J). The following screen shows and analysis is started by clicking on the '*Process*' button (K).

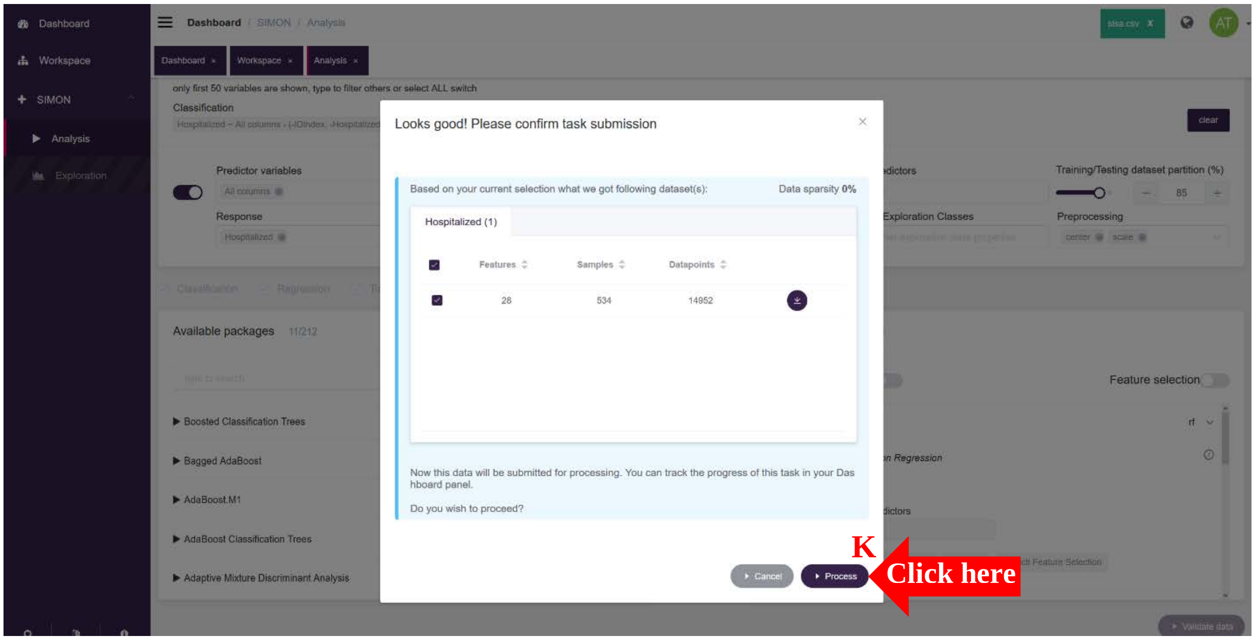


New window '*Dashboard*' will open immediately and SISA analysis will be created and initiated (L). To assess the current status of the analysis, we look under '*Status*' column (M), '*Processing*' means that the analysis is going, once analysis is complete it says '*Completed*' in green. '*Sparsity (%)*' column tells us the percentage of missing values in the dataset; ‘*Models processed*’, number of processed models; ‘*Successful models*’, number of successfully finished models. In the ‘*Operations*’ column users can get information about the models and their performance measures during the analysis (first button, “circled *i*” icon), download dataset (second full green button) and delete the analysis (third red button).

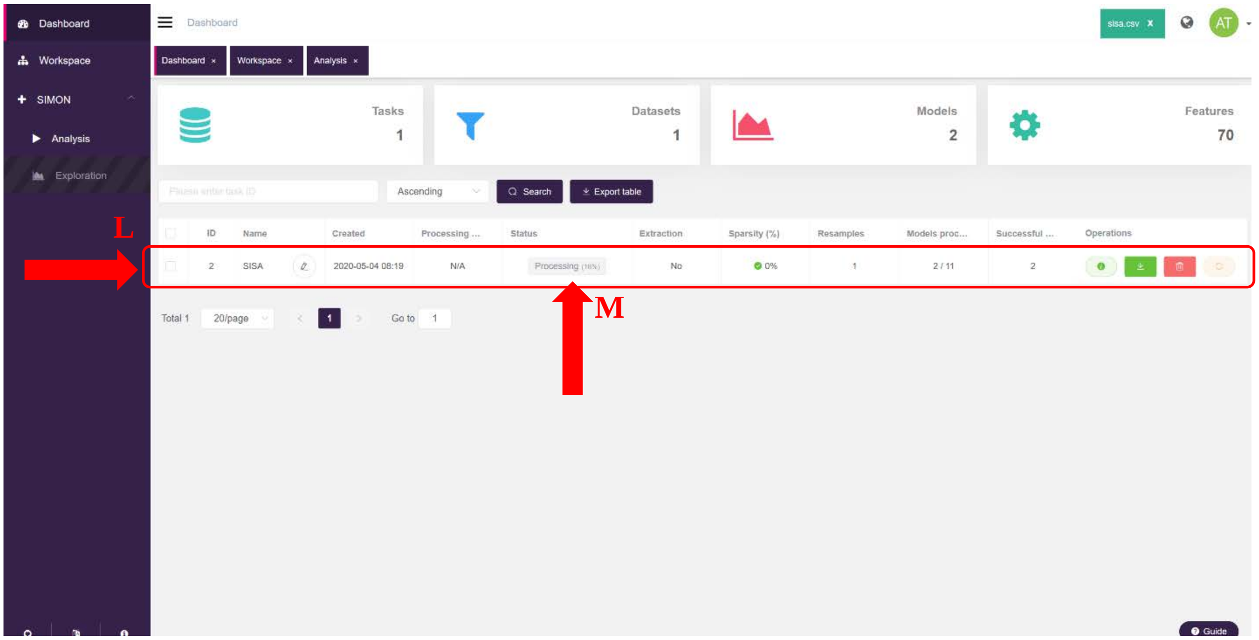


**Step 4. Model evaluation and selection.** After the analysis is done, users can explore built predictive models by clicking on the checkbox next to the SISA analysis row (A). Upon selection of SISA analysis row, the ‘*Exploration*’ tab (B) becomes available in the menu on the left-hand side. By clicking on the ‘*Exploration*’ tab, new window opens where users can explore built models.

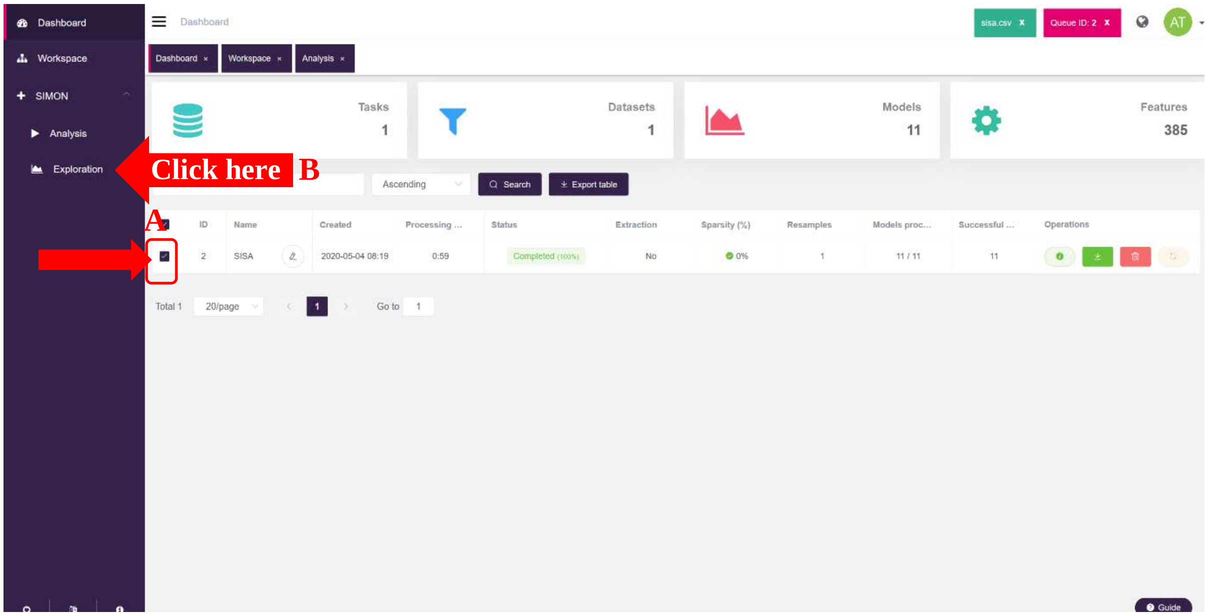

Now, in the new window, users select desired performance measurements by clicking on the drop-down menu in the input Box 1. SIMON calculates different performance measurements for training set (train AUC, train F1, train prAUC, train recall, train precision, train sensitivity and train specificity) and test set (accuracy, F1, kappa, predict AUC, predict prAUC, precision, recall, sensitivity and specificity). For the SISA dataset, we choose AUROC as performance measurement by selecting train AUC and predict AUC. Now, we click in the dataset in the Box 2 (C). This opens Box 3 containing table of all models that were built. We order models based on the train AUC value (D). The model built with the highest train AUC was built with the *sda* algorithm. To compare models, users select desired models by clicking the check box next to each model. We will compare top five models by clicking in the 'select all' check box (E). Once all five models are selected, users can download the table with all models and performance measurements as CSV file and models with the data as RData objects by clicking on F. The initial dataset, training and test set can be saved as CSV files by clicking the download icon next to the dataset row in the Box 2 (G).

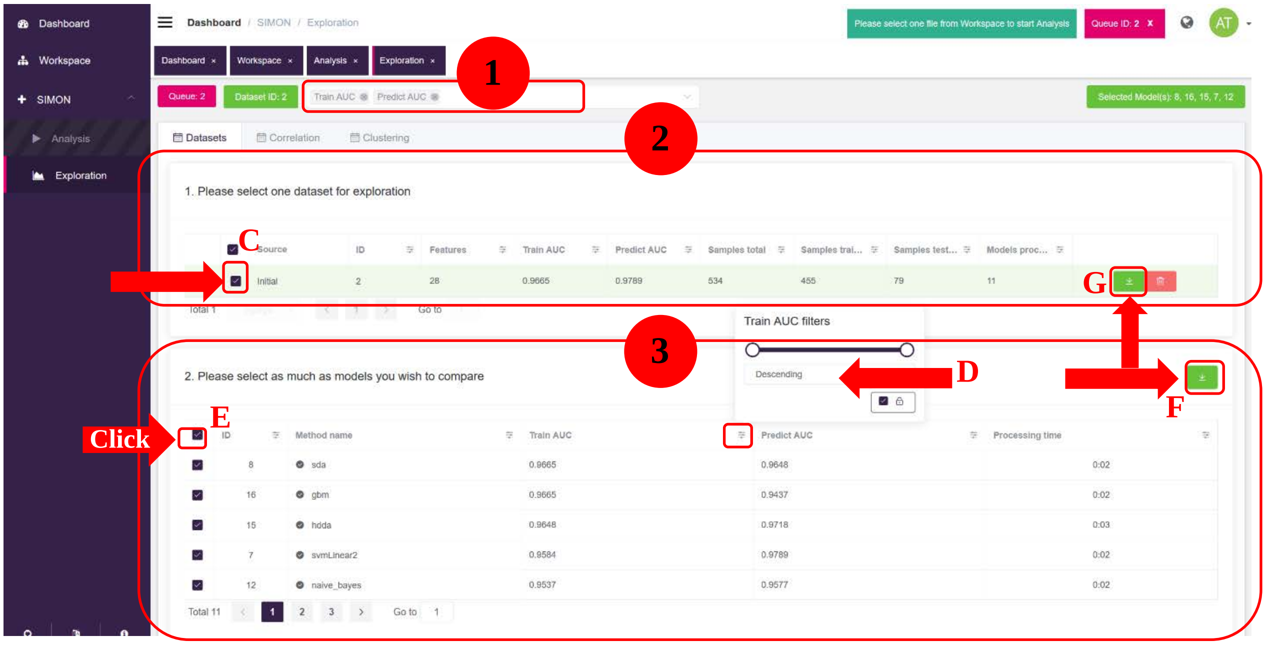


By selecting models to compare, the '*Training Summary*' tab will appear below Box 3 (H). Users must select at least two models for the tab to appear. Here, users visualize model comparison and can download box plots graphs showing performance measures calculated for the training set (I) and ROC plots for the training set (J) for all models selected as SVG files. To select all 11 models, as we did in the Figure 1, users must navigate to pages 2 and 3 and click select all check box.

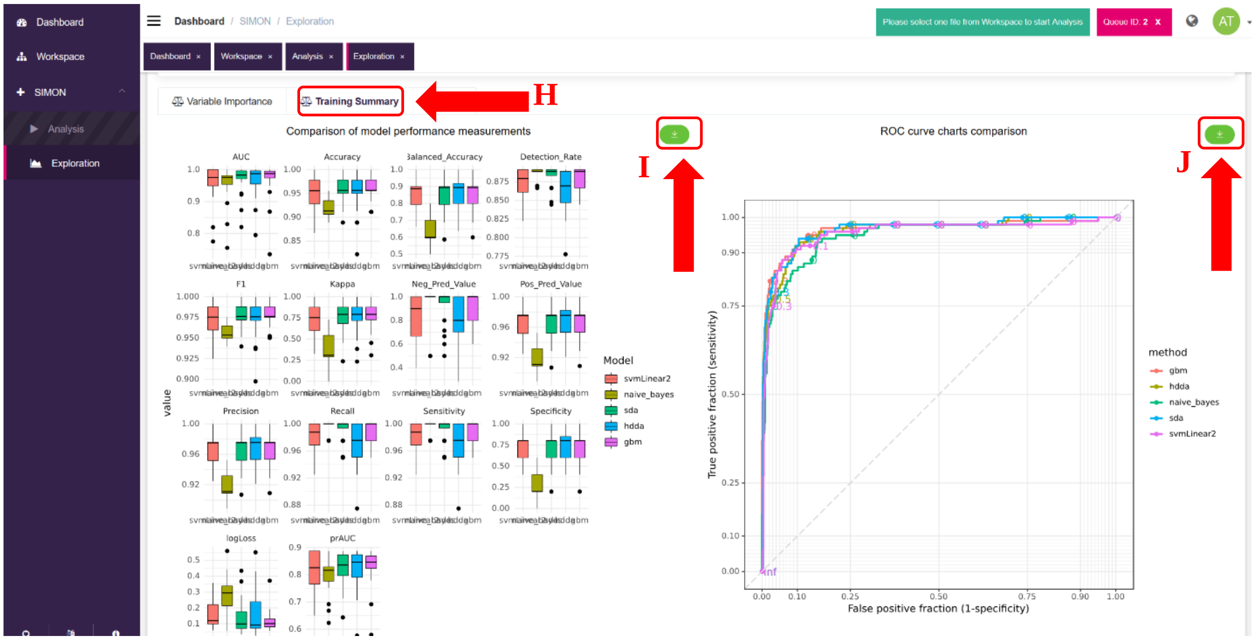


**Step 5. Feature selection.** To explore features that contributed the most to the sda model, we select only sda model in the Box 3 and click the '*Variable importance*' tab next to the *Training Summary*' tab (A). This opens table where features are ranked based on the Variable Importance Score ('*Score*' column). Features that have Variable Importance score above 50 are highlighted in green (B). The table can be downloaded as the CSV file by clicking download button (C). Users can select two or more models and compare ranking of features across models.

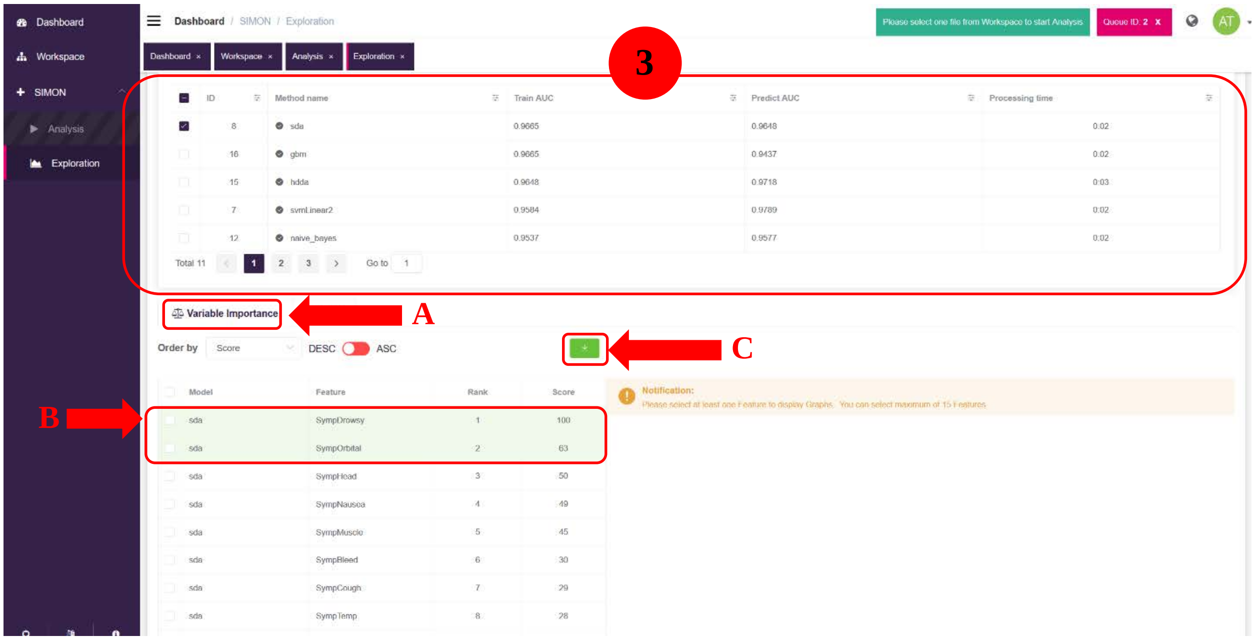

By selecting desired features, users can visualize distribution of data between both groups using dot plots. Plots can be adjusted by selecting '*Theme*', '*Color*', font size ('*Font*'), dot size ('*Dot*') and height/width ratio ('*Ratio*') as described in the ggplot2 R package (https://ggplot2.tidyverse.org/) (D). To apply changes to the graphs users must press '*Redraw plot*' red button (E) and graphs can be downloaded as SVG files by pressing download button (F).

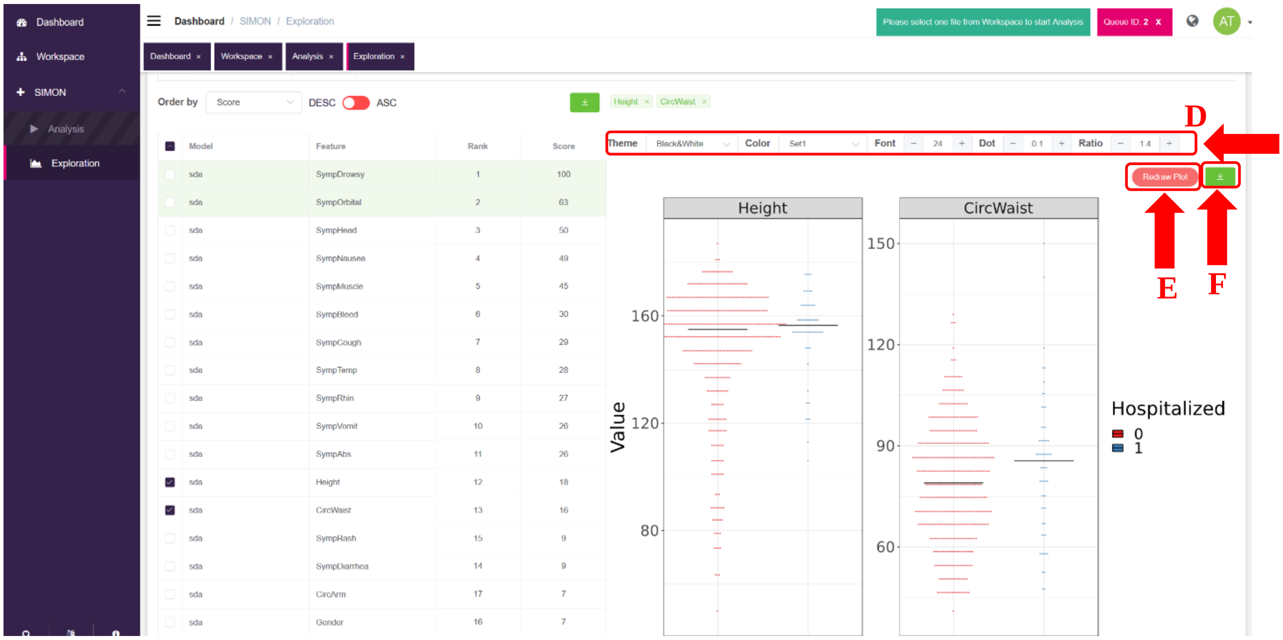


**Step 6. Exploratory analysis.** In the 'Exploration' window, upon selection of dataset for analysis, two tabs are shown: '*Correlation*' and '*Clustering*' (A). By clicking on '*Correlation*' tab, users can perform correlation analysis on the selected dataset using three different correlation methods (Pearson, Kendall and Spearman) and different parameters can be applied by clicking the '*Plot image*' red button (B). Correlation plot can be saved as SVG file by clicking download button (C). '*Clustering*' tab will be explained in the Use case 2.

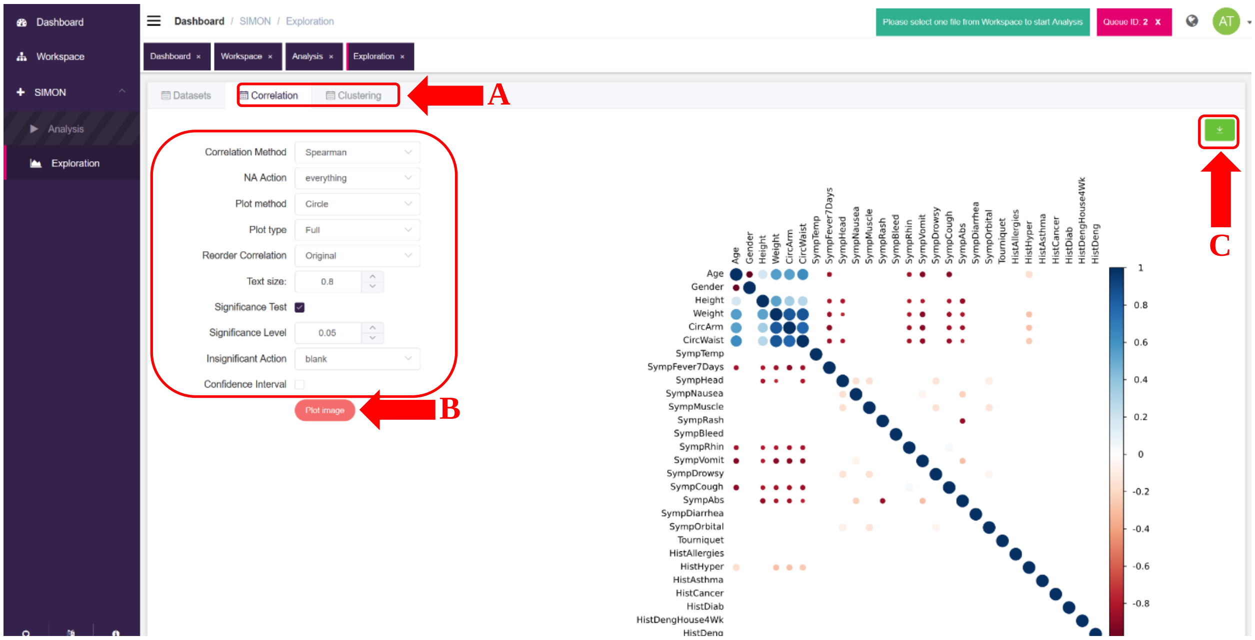


**Use case 2. Predicting antibody signature to mediate protection against *Salmonella* Typhi challenge infection.**

The first two steps (Step 1. Uploading data and Step 2. Selecting dataset.) are the same as explained under Use case 1, therefore we will start from Step 3.

**Step 3. Setting-up analysis.** To perform SIMON analysis on the VAST dataset, users must select all predictor variables by clicking the button next to the *'Predictor variables'* input form (A) and *'Diagnosis'* column as the outcome in the *'Response'* input form (B). *'Vaccine'* column is selected under *'Additional Exploration Classes'* (C). The initial dataset is split into training (75% of the data) and test sets (25% of the data) (D) and we applied 'center' and 'scale' as pre-processing steps (E). In total, five ML algorithms were selected (F). Since the VAST dataset has missing values, in the first step of SIMON we will use *'Multi-set intersection'* function (G).

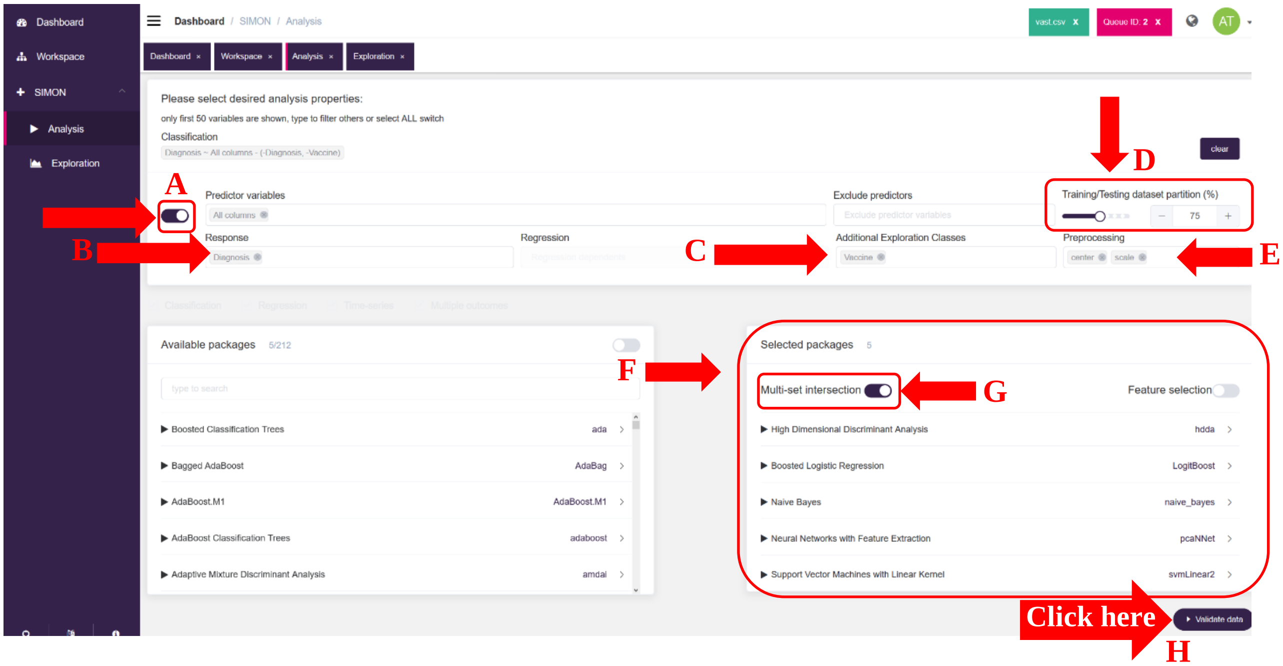


By clicking ‘*Validate data’* button (H), multi-set intersection function will generate resamples and the pop-up window shows 206 generated resamples with different number of ‘Features’ and donors (‘Samples’ column). Each resample can be saved by clicking on the download button and analysis can be performed by selected resamples. In the VAST dataset, we performed analysis on 58 resamples with 40 or more samples in total. Click *‘Process’* button to start analysis.

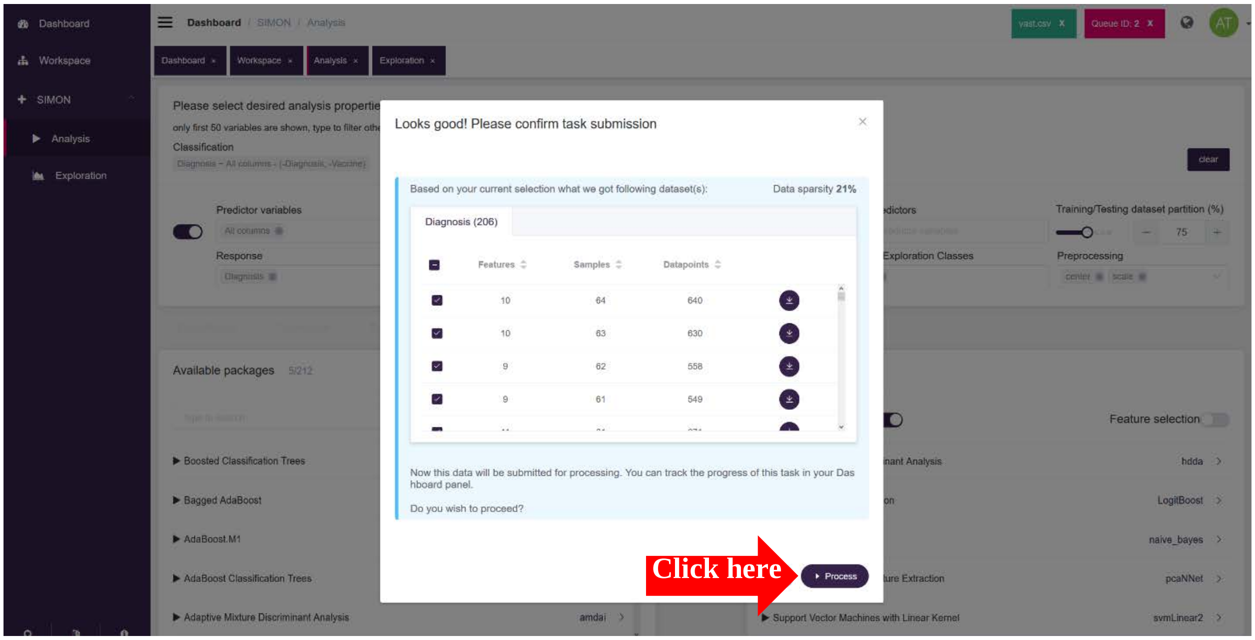


**Step 4. Model evaluation and selection.** We open *‘Exploration’* window (tab becomes available upon selection of VAST analysis row in the *‘Dashboard’*) and select train AUC and predict AUC as performance

measurements (A). Then, we order datasets (i.e. resamples) based on the maximum train AUC value and using slider remove all models with train AUC value below 0.72 (B). Now, we select the first dataset with 10 'Features' (C) and 45 'Samples total' (D) and in the table below, we explore all models built for that dataset by ordering models based on the train AUC (E). The model built with the highest performace measurements was built with the pcaNNet algorithm (F), but despite high train AUC value, that model does not perform so well on the unseen test data (predict AUC 0.64). Such a model is considered overfitted. We must examine other resamples to find optimal models with better perfomance on the test data. Each dataset/resample and accompanying split into training and test sets can be saved as CSV file by clicking the download button (G). To highest perfoming model was built on the dataset/resample that has 13 features and 47 donors (samples) (ID 38). We select that dataset and evaluate models using box plots and ROC curves as described for Use case 1.

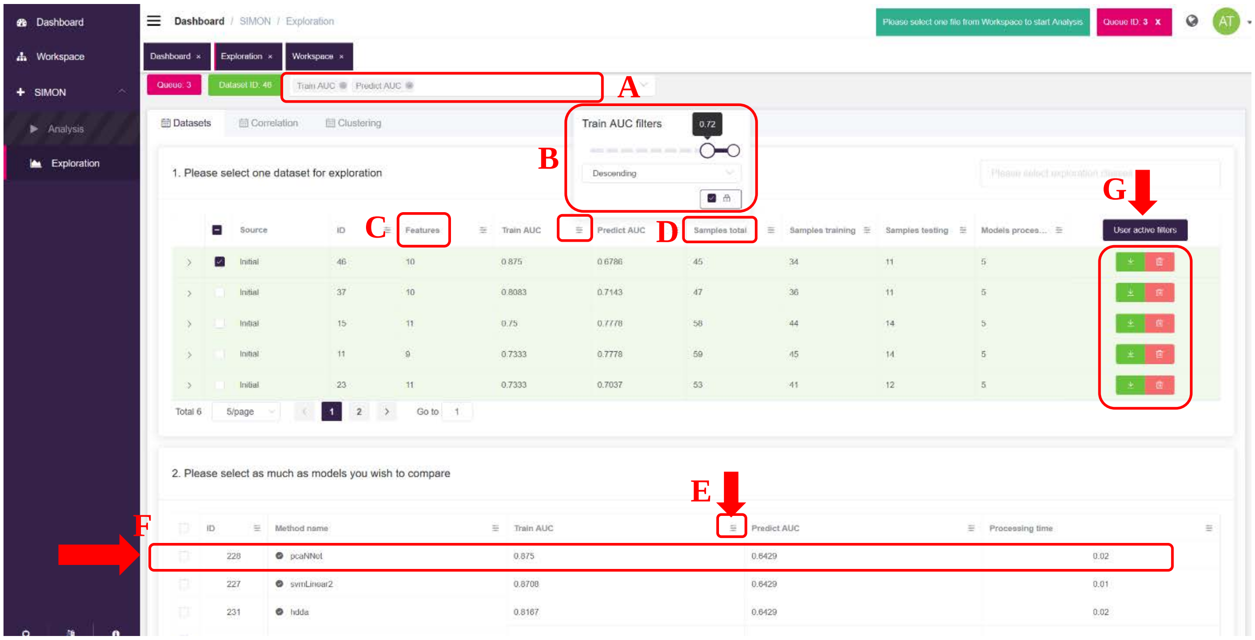


**Step 5. Feature selection.** Upon selecting the best performing model built with the naïve Bayes algorithm, we can explore the features that contributed the most to this model in the '*Variable importance*' tab. The variable importance score table can be downloaded as a CSV file and graphs as SVG files by clicking download buttons (A and B).

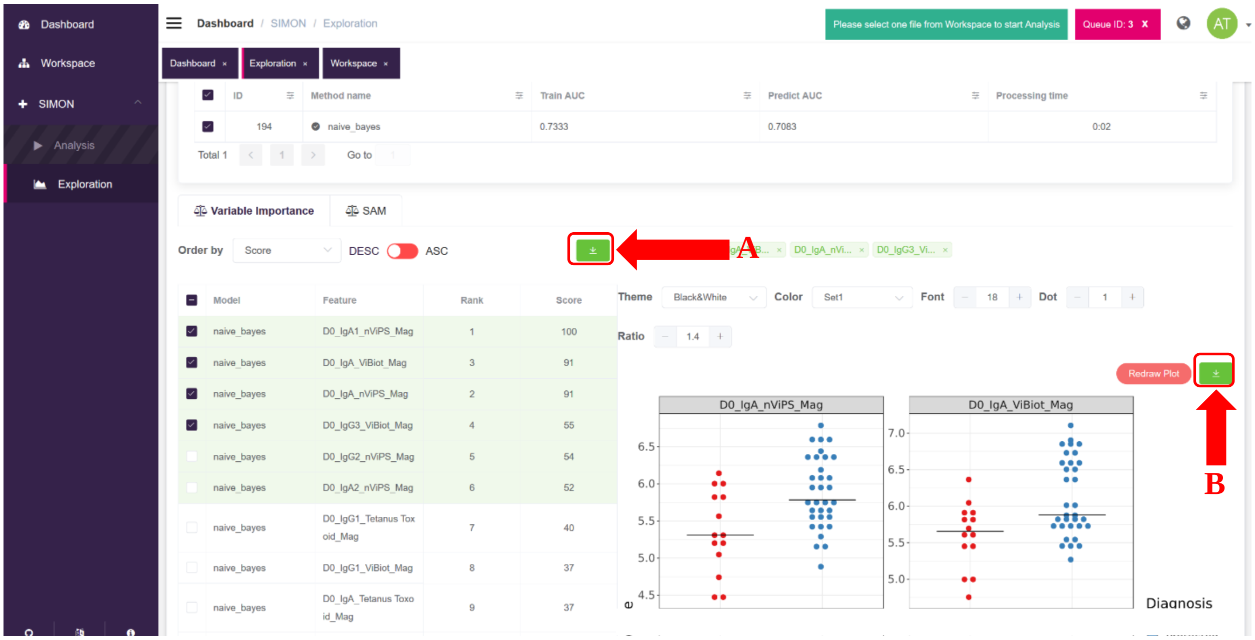


**Step 6. Exploratory analysis.** In addition to the *'Correlation'* tab explained above in the Use case 1, users can perform clustering analysis in the *'Clustering'* tab. In the VAST dataset we want to explore if the individuals are grouped based on the vaccine they received ('*Vaccine*' column selected under *'Additional Exploration Classes'*). We select *'Diagnosis'* and *'Vaccine'* as columns and 3 top features as rows. After setting up the desired parameters for the clustering analysis, we click *'Plot image'* button (A). The heatmap can be saved as a CSV file by clicking on the download button (B). We can also perform clustering analysis as described above in Use case 1.

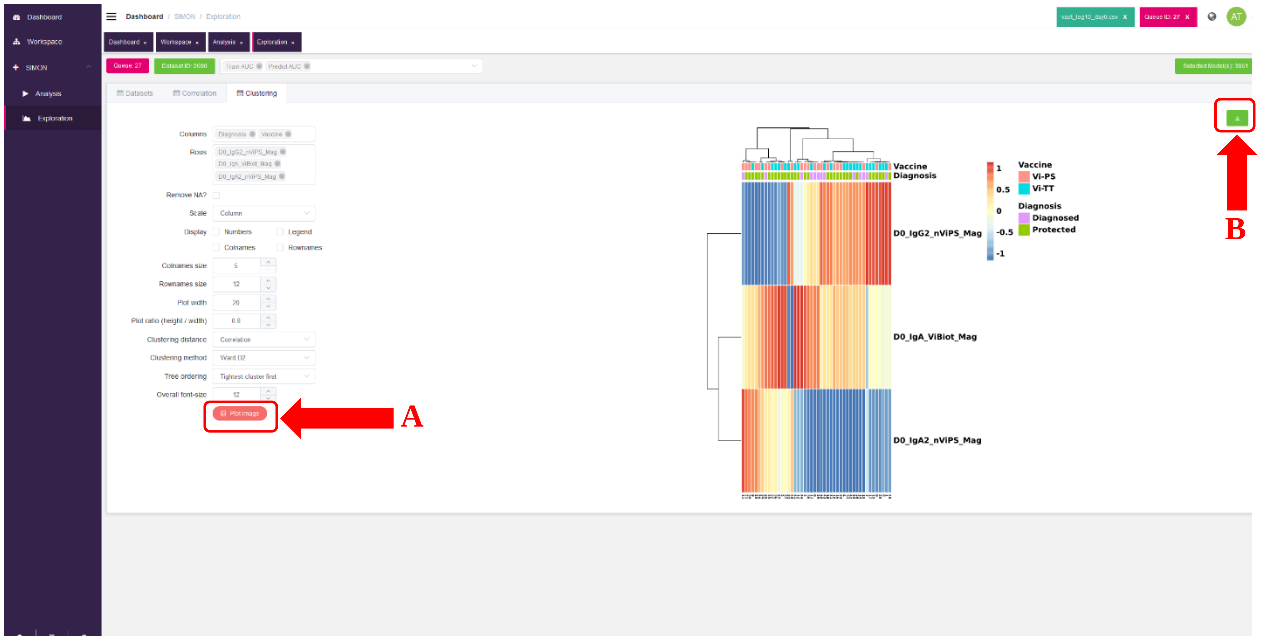

**Use case 3. Identifying cellular immune signature associated with high-level of physical activity.**

Steps 1-2 and 4-6 were performed as described above for the first two use cases.

**Step 3. Setting-up analysis.** For the Cyclists dataset, we used all columns as *'Predictors variables'* (A) and outcome column as the *'Response'*(B). Other parameters, training/test split and preprocessing were performed as shown in the screenshot below. Similar to the use case 2, we used multi-set intersection function for the initial dataset to find resamples (C). In total, 146 resamples were identified and analysis was performed using all resamples.

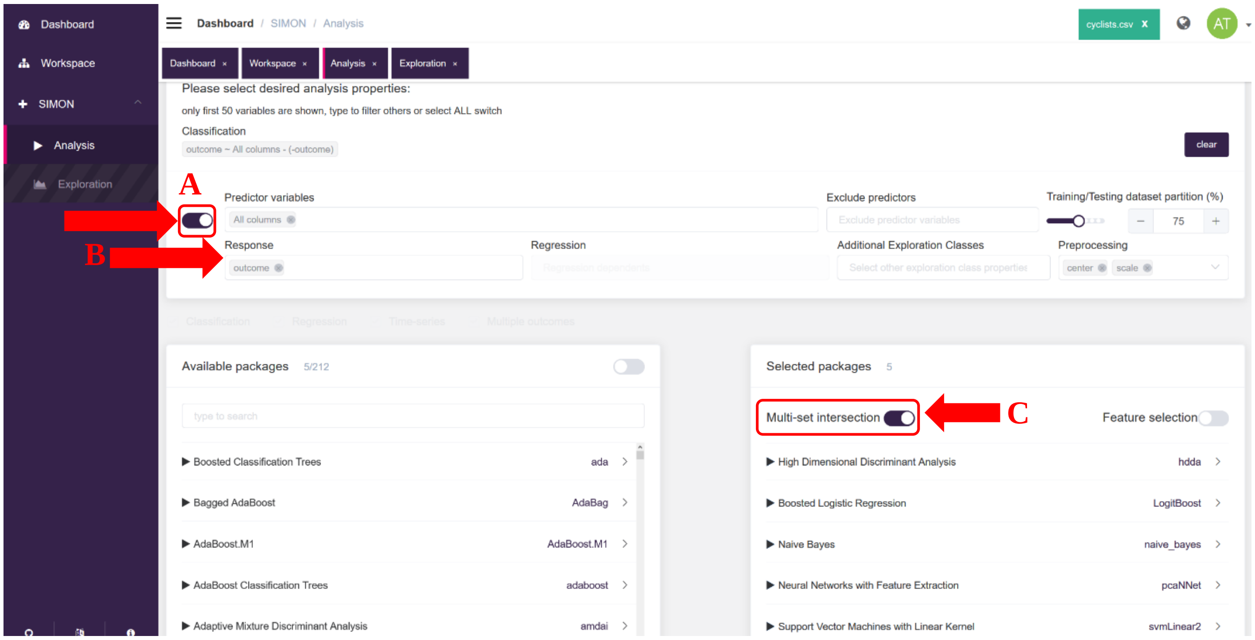


**Use case 4. Building predictive model for the early-stage detection of colorectal cancer using microbiome.**

Steps 1-2 and 4-6 were performed as described above.

**Step 3. Setting-up analysis.** After uploading and selecting the Zeller dataset, in the *'Analysis'* window, we selected all columns as *'Predictors variables'* (A) and we typed *'Study condition'* in the *'Response'* input form to find the outcome column (B). The initial dataset was divided 75% into training and 25% test set (C). For the preprocessing we applied '*center*', '*scale'* and remove near zero-variance ('*nzv*') (D). In total, five ML algorithms were selected (E) and analysis was started by pressing '*Validate data'* button.

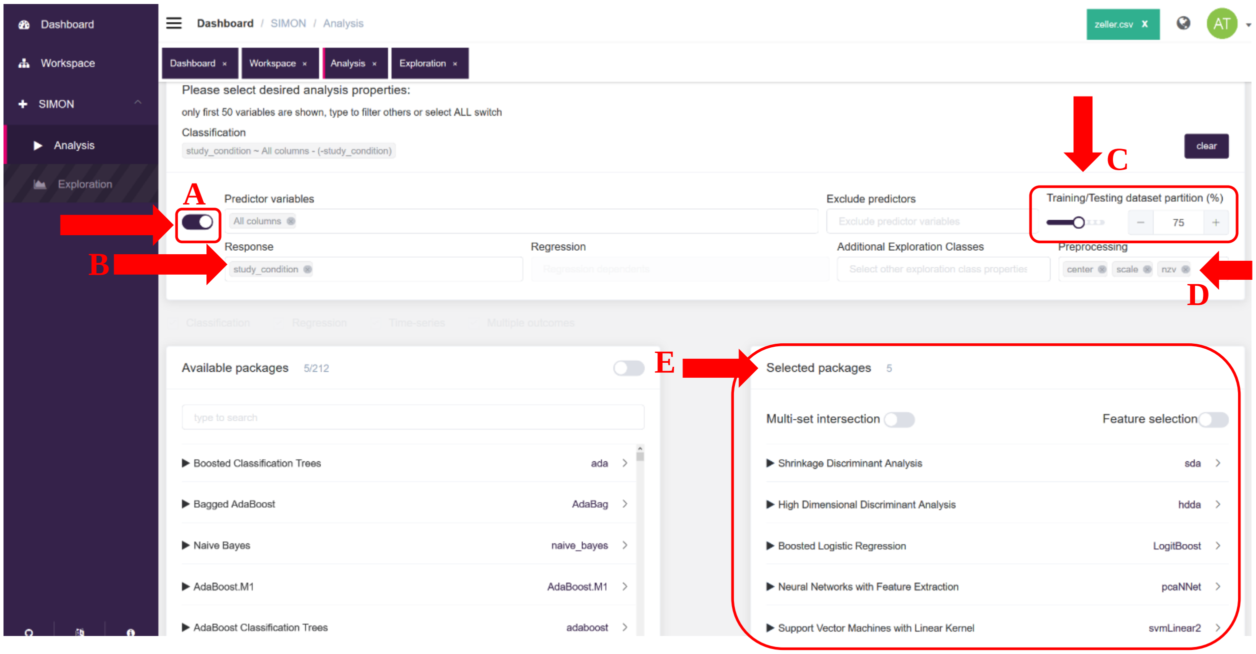


**Use case 5. Building predictive model for detection of liver hepatocellular carcinoma cells using transcriptome data.**

Steps 1-2 and 5-6 were performed as described above for the other Use cases.

**Step 3. Setting-up analysis. For the LIHC dataset, analysis was started with** selecting all columns as *'Predictors variables'* (A) and *'Group'* column (tumor or healthy cells) as the *'Response'* (B). The initial dataset was divided 75% into training and 25% test set (C). For the preprocessing we applied '*center*', '*scale*' and remove near zero-variance ('*nzv*') (D). In total, five ML algorithms were selected (E) and analysis was started ('*Validate data*' button).

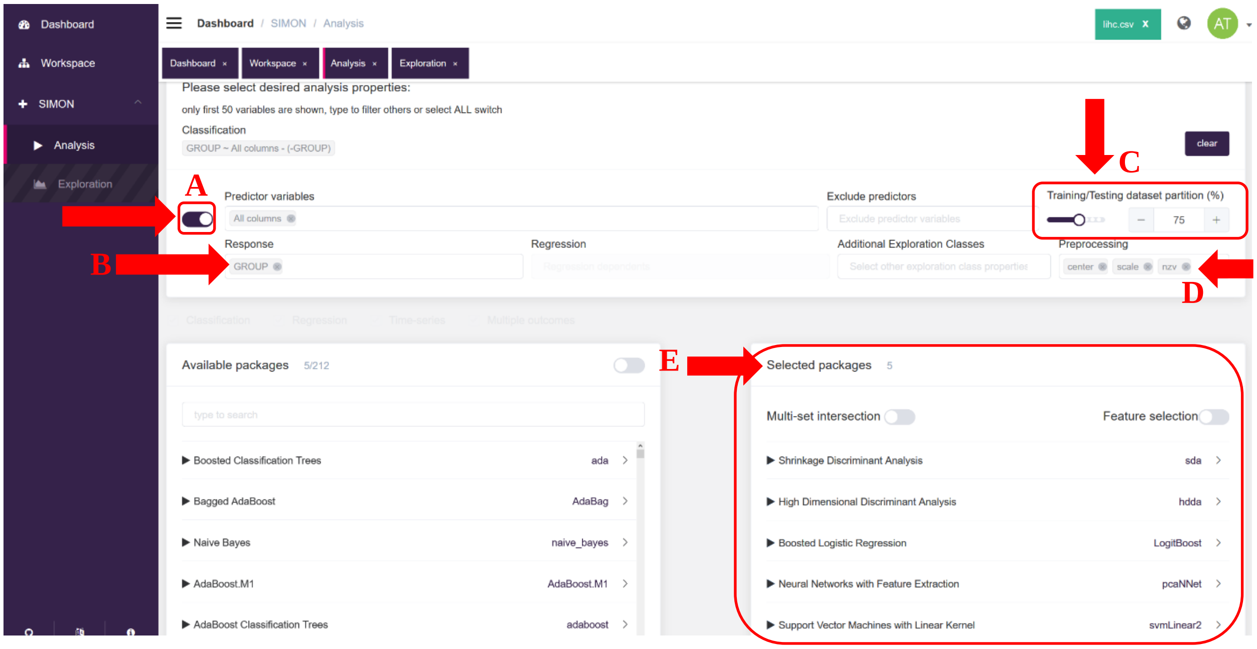


**Step 4. Model evaluation and selection.** The LIHC is the example of highly imballanced dataset, therefore in the '*Exploration*' window (tab becomes available upon selection of LIHC analysis row in the '*Dashboard*') and select precision-recall AUC (train prAUC for the training set and prAUC for the test set) (A). The models are then ranked based on the train prAUC (B). The first model that has high train prAUC value, also performed well on the left-out test set. We save the generated model by clicking the download button (C). Visualization of model performance measurements is performed as described for other Use cases.

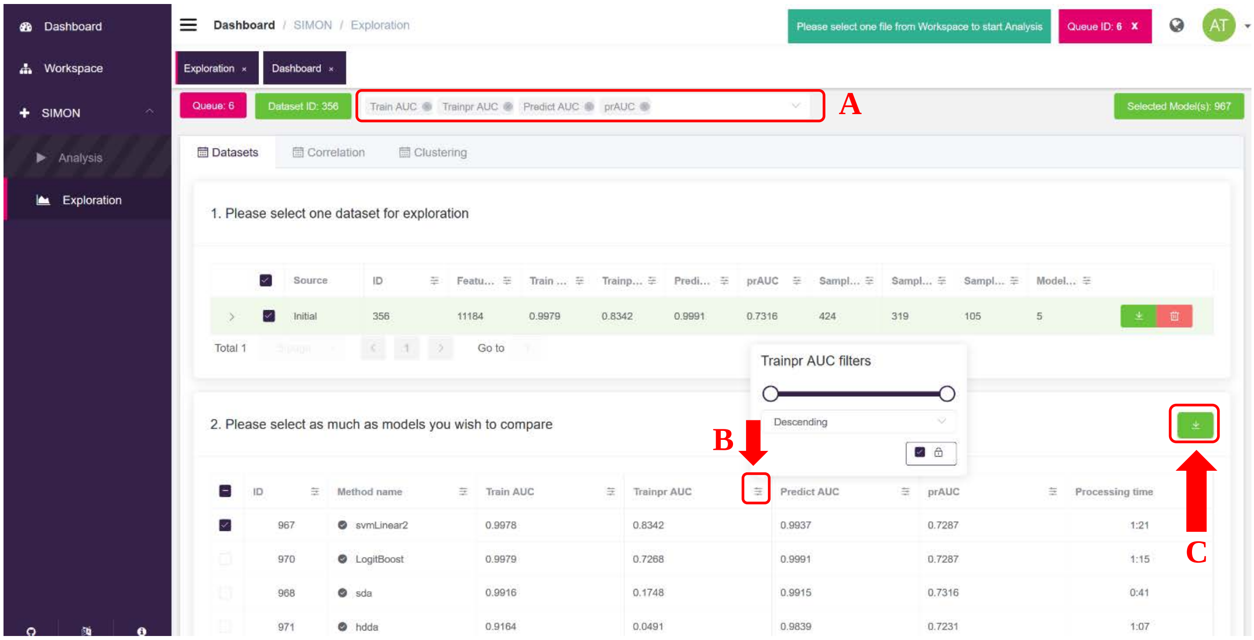

#### Online Methods References

##### List of references of R packages used for Supplementary Table 1:

32. Package: gam - Trevor Hastie (2017). gam: Generalized Additive Models. R package version 1.14-4. https://CRAN.R-project.org/package=gam
33. Package: h2o - Erin LeDell, Navdeep Gill, Spencer Aiello, Anqi Fu, Arno Candel, Cliff Click, Tom Kraljevic, Tomas Nykodym, Patrick Aboyoun, Michal Kurka and Michal Malohlava (2020). h2o: R Interface for the 'H2O' Scalable Machine Learning Platform. R package version 3.28.0.4. https://CRAN.R-project.org/package=h2o
34. Package: gbm - Brandon Greenwell, Bradley Boehmke, Jay Cunningham and GBM Developers (2019). gbm: Generalized Boosted Regression Models. R package version 2.1.5. https://CRAN.R-project.org/package=gbm
35. Package: glm2 - Ian C. Marschner (2011). Fitting generalized linear models with convergence problems. The R Journal, 3(2), 12-15. URL https://CRAN.R-project.org/package=glm2
36. Package: MASS - Venables, W. N. & Ripley, B. D. (2002) Modern Applied Statistics with S. Fourth Edition. Springer, New York. ISBN 0-387-95457-0
37. Package: gpls - Beiying Ding (2018). gpls: Classification using generalized partial least squares. R package version 1.54.0.
38. Package: hda - Gero Szepannek (2016). hda: Heteroscedastic Discriminant Analysis. R package version 0.2-14. https://CRAN.R-project.org/package=hda
39. Package: HDclassif - Laurent Berge, Charles Bouveyron, Stephane Girard (2012). HDclassif: An R Package for Model-Based Clustering and Discriminant Analysis of High-Dimensional Data. Journal of Statistical Software, 46(6), 1-29. URL http://www.jstatsoft.org/v46/i06/
40. Package: pls - Bjørn-Helge Mevik, Ron Wehrens and Kristian Hovde Liland (2019). pls: Partial Least Squares and Principal Component Regression. R package version 2.7-2. https://CRAN.R-project.org/package=pls
41. Package: kknn - Klaus Schliep and Klaus Hechenbichler (2016). kknn: Weighted k-Nearest Neighbors. R package version 1.3.1. https://CRAN.R-project.org/package=kknn
42. Package: klaR - Weihs, C., Ligges, U., Luebke, K. and Raabe, N. (2005). klaR Analyzing German Business Cycles. In Baier, D., Decker, R. and Schmidt-Thieme, L. (eds.). Data Analysis and Decision Support, 335-343, Springer-Verlag, Berlin.

43. Package: logicFS - Holger Schwender and Tobias Tietz (2018). logicFS: Identification of SNP Interactions. R package version 2.2.0.
44. Package: caTools - Jarek Tuszynski (2020). caTools: Tools: moving window statistics, GIF, Base64, ROC, AUC, etc. R package version 1.17.1.4. https://CRAN.R-project.org/package=caTools
45. Package: LogicReg - Charles Kooperberg and Ingo Ruczinski (2019). LogicReg: Logic Regression. R package version 1.6.2. https://CRAN.R-project.org/package=LogicReg
46. Package: class - Venables, W. N. & Ripley, B. D. (2002) Modern Applied Statistics with S. Fourth Edition. Springer, New York. ISBN 0-387-95457-0
47. Package: mda - S original by Trevor Hastie & Robert Tibshirani. Original R port by Friedrich Leisch, Kurt Hornik and Brian D. Ripley. (2017). mda: Mixture and Flexible Discriminant Analysis. R package version 0.4-10. https://CRAN.R-project.org/package=mda
48. Package: HiDimDA - Antonio Pedro Duarte Silva (2015). HiDimDA: High Dimensional Discriminant Analysis. R package version 0.2-4. https://CRAN.R-project.org/package=HiDimDA
49. Package: RSNNS - Christoph Bergmeir, Jose M. Benitez (2012). Neural Networks in R Using the Stuttgart Neural Network Simulator: RSNNS. Journal of Statistical Software, 46(7), 1-26. URL http://www.jstatsoft.org/v46/i07/
50. Package: keras - JJ Allaire and François Chollet (2019). keras: R Interface to 'Keras'. R package version 2.2.5.0. https://CRAN.R-project.org/package=keras
51. Package: FCNN4R – G Klima (2016), FCNN4R: Fast Compressed Neural Networks for R, R package version 0.6.2.
52. Package: monmlp - Alex J. Cannon (2017). monmlp: Monotone Multi-Layer Perceptron Neural Network. R package version 1.1.4. https://CRAN.R-project.org/package=monmlp
53. Package: nnet - Venables, W. N. & Ripley, B. D. (2002) Modern Applied Statistics with S. Fourth Edition. Springer, New York. ISBN 0-387-95457-0
54. Package: naivebayes - Michal Majka (2017). naivebayes: High Performance Implementation of the Naive Bayes Algorithm. R package version 0.9.1. https://CRAN.R-project.org/package=naivebayes

77. Package: rpart - Terry Therneau, Beth Atkinson and Brian Ripley (2017). rpart: Recursive Partitioning and Regression Trees. R package version 4.1-11. https://CRAN.R-project.org/package=rpart
78. Package: rpartScore - Giuliano Galimberti, Gabriele Soffritti and Matteo Di Maso (2012). Classification Trees for Ordinal Responses in R: The rpartScore Package. Journal of Statistical Software, 47(10), 1-25. URL http://www.jstatsoft.org/v47/i10/
79. Package: RRF - H. Deng (2013). Guided Random Forest in the RRF Package. arXiv:1306.0237.
80. Package: sda - Miika Ahdesmaki, Verena Zuber, Sebastian Gibb and Korbinian Strimmer (2015). sda: Shrinkage Discriminant Analysis and CAT Score Variable Selection. R package version 1.3.7. https://CRAN.R-project.org/package=sda
81. Package: sdwd - Boxiang Wang and Hui Zou (2020). sdwd: Sparse Distance Weighted Discrimination. R package version 1.0.3. https://CRAN.R-project.org/package=sdwd
82. Package: ipred - Andrea Peters and Torsten Hothorn (2019). ipred: Improved Predictors. R package version 0.9-9. https://CRAN.R-project.org/package=ipred
83. Package: sparseLDA - Line Clemmensen and contributions by Max Kuhn (2016). sparseLDA: Sparse Discriminant Analysis. R package version 0.1-9. https://CRAN.R-project.org/package=sparseLDA
84. Package: spls - Dongjun Chung, Hyonho Chun and Sunduz Keles (2019). spls: Sparse Partial Least Squares (SPLS) Regression and Classification. R package version 2.2-3. https://CRAN.R-project.org/package=spls
85. Package: vbmp - Nicola Lama and Mark Girolami (2018). vbmp: Variational Bayesian Multinomial Probit Regression. R package version 1.50.0. http://bioinformatics.oxfordjournals.org/cgi/content/short/btm535v1
86. Package: VGAM - Thomas W. Yee (2015). Vector Generalized Linear and Additive Models: With an Implementation in R. New York, USA: Springer.
87. Package: wsrf - He Zhao, Graham J. Williams, Joshua Zhexue Huang (2017). wsrf: An R Package for Classification with Scalable Weighted Subspace Random Forests. Journal of Statistical Software, 77(3), 1-30. doi:10.18637/jss.v077.i03
88. Package: xgboost - Tianqi Chen, Tong He, Michael Benesty, Vadim Khotilovich, Yuan Tang, Hyunsu Cho, Kailong Chen, Rory Mitchell, Ignacio Cano, Tianyi Zhou, Mu Li,

Junyuan Xie, Min Lin, Yifeng Geng and Yutian Li (2020). xgboost: Extreme Gradient Boosting. R package version 1.0.0.2. https://CRAN.R-project.org/package=xgboost
